# Epigenetic perturbation unravels diverse origins, state conversions and de-differentiation of cortical astrocytes

**DOI:** 10.64898/2026.09.11.750559

**Authors:** Maira Mirza, Tobias Hohl, Camila L. Fullio, Camila Vidos, Johan Rollin, Thomas Manke, Tanja Vogel

## Abstract

The role of epigenetic modifiers in astrogenesis is less understood compared to neurogenesis. For example, the Disruptor of telomeric silencing 1 like, DOT1L, conferring H3K79 methylation safeguards neural progenitors from premature differentiation in various brain regions, however, its role in astrogenesis has not yet been reported.

Here, we decipher its role during astrogenesis using an *Emx1-cre* driven DOT1L conditional knock-out during the development of mouse cerebral cortex from E12.5 till P0. We describe two astrocyte types at P0, distinguishable by expression of phagocytosis markers genes. DOT1L loss-of-function (LOF) increased the number of those astrocytes that express fewer phagocytosis marker genes, while it affects a larger number of differentially expressed genes in those astrocytes that strongly express homeostasis marker genes. Our findings establish DOT1L as a barrier between these astrocyte states, limiting differentiation of the astrocytes expressing phagocytosis marker genes.

We also identify multiple developmental origins of astrocytes. In addition to apical progenitors (AP), the prevailing stem cell population considered as origin of cortical astrocytes, we identify neural stem cells (NSC) in the dorsal telencephalon differentiating towards astrocytes, and *Nkx2.1*-lineage derived astrocytes, originating from the ventral telencephalon. This finding refines the prevailing view that dorsal telencephalon APs are the only source of embryonic cortical astrocytes, and enter astrogliogenesis after neurogenesis. We show that the expression of *Emx1* discriminates different astrocyte origins during cortical development, and we describe a new set of transcription factors (TFs) that mark the state conversion between the homeostasis or phagocytosis expression programs in astrocytes. DOT1L orchestrates different TF networks, of which *Lhx2, Nfia, Sox6*, and *Creb5* drive astrogliogenesis, and *Nfia, Nfib*, and *Tcf4* affect astrocyte state conversion. Importantly, our analysis also shows that NSCs, APs and astrocytes de-differentiate within this lineage trajectory, and lowering the DOT1L-mediated epigenetic barrier strongly favours appearance of astrocytes with limited phagocytosis properties and reactivation of progenitor programs.

## Introduction

Brain development depends crucially on orchestrated gene transcription to build functional entities, in which the activities of neurons, astrocytes, or oligodendroglia are tightly coordinated. These different cell types arise in more or less sequential order with the progression of time, seemingly from the same or similar pools of stem cells [1]. For example, in the current understanding of mouse cerebral cortex development, apical and basal progenitors (AP, BP; synonymously radial glia cells (RGC) or intermediate progenitor cells (IPC)) produce mainly glutamatergic neurons during prenatal development, switch to astrogliogenesis at late embryonic stages [2], and produce oligodendroglia postnatally [3]. Despite prenatal origins, the majority of astrocytes arise postnatally, through local amplification [4]. However, the developmental origin of astrocytes residing in the cerebral cortex from a common, earlier neurogenic ancestor is a prevailing view, which is challenged by recent findings. For example, multiclonal lineage tracing showed astrocyte origins from embryonal and postnatal progenitors [5]. Astrocytes not only appear at different developmental time points [6], but also in specific CNS regions, for example, in discrete layers of the cerebral cortex [7,8]. The heterogeneity of astrocyte origins is further supported by the finding of a dual origin of astrocytes from either *Emx1*- or *Olig2*-expressing progenitor [9]. In addition, ventral progenitors, which are the main source for cortical GABAergic interneurons, also generate astrocytes that settle in the cortical pallium [10,11]. Consequently, models emerge that attribute different astrocyte subtypes to specialized functions [5], one of which is synapse formation [9].

In functional terms, astrocytes interact with other cell types, which is essential for proper neuronal communication. Neuronal activity relies on precise homeostatic conditions including nutrition, ion availability, or transmitter turnover, all of which are under the control of astrocytes. Furthermore, astrocytes influence brain homeostasis by interacting with blood vessels, but also by exerting phagocytosis. They are also majorly involved in brain repair processes. In this context, they become reactive, and in this state their appearance is a hallmark of inflammatory or degenerative processes.

Since astrocytes keep their proliferative capacities and are attracted to locations of acute brain injury, they are a given target for reprogramming attempts, with the aim to replace, for example, degenerated neurons. During astrocyte reprogramming, neuronal programs are generally reactivated for example through expression of *Ngn2* [12]. Another possibility is to reprogram astrocytes to *Ascl1*-expressing progenitor states, through reactivation of either a gene set conferring stem cell character, i.e. *Oct4, Nanog*, and *Sox2* [13], or, alternatively, *Dlx2* [14], *Foxg1, Sox2*, or *Brn2* [15] expression, which reprogram astrocytes to neural progenitor cells. These findings indicate the plasticity of astrocytes to convert into other cell types.

Clearly, all aspects of astrocyte generation, network integration, functional specification as well as their plasticity to adapt different functional states or revert to neurons or neural progenitors, depends on transcriptional programs and tight control thereof. In this context, diverse transcription factors (TF) have been highlighted, which are implicated generally in brain development, and impinge on balancing neuro- or astrogliogenesis. Examples of instrumental TF for astrogenesis in the cerebral cortex include *Nfia, Sox9, Stat3, Lhx2*, or *Foxg1* [16-19]. Moreover, *Tcf4* suppresses the astrocyte fate of oligodendroglia progenitors originating from ventral progenitor populations [20].

Nowadays, the regulative layer of epigenetic modifications receives attention for its impact on developmental processes, where it coordinates activity in regard to a higher-order adaptation of the chromatin towards accessibility of transcriptional executors. However, little data are available as of yet that expose a role for epigenetic regulation of astrogliogenesis, astrocyte specification and function, during inflammatory processes or reprogramming [21-23]. Altered DNA methylation at STAT-binding sites is one prominent example of how epigenetic processes regulate the transition from neuro-to astrogenesis [19]. Further work exposed altered epigenetic landscapes of astrocytes from cortical and cerebellar regions in the postnatal mouse brain [24]. But generally, we are trailing behind in understanding the epigenetic regulation of astro-compared to neurogenesis in pre- and early postnatal development.

Recent studies demonstrated critical roles of epigenetic regulation for neurogenesis in the developing cerebral cortex, among which is the histone H3 lysine 79 (H3K79) methylation mediated by the disruptor of telomeric silencing 1 like, DOT1L [25-28]. DOT1L has a gate-keeper function for cell transition during differentiation, as loss or inhibition of DOT1L during neurogenesis increases the production of neurons at the expense of the progenitor pool. As neurogenesis precedes astrogliogenesis, it is as of yet unclear whether genetic deletion of DOT1L affects the ability of cortical progenitors to generate astrocytes, specific subtypes or activity states.

Here, we report on conditional *Dot1l* deletions in mice, which we studied at different developmental points at single cell level. Our data indicate that DOT1L loss-of-function (LOF) through *Emx1-cre* (1) drives astrocyte state-specific expression programs, (2) regulates conversion between astrocyte states, diverging by gene expression related to phagocytosis, (3) limits de-differentiation of astrocytes to progenitor states, and (4) regulates a TF network, driving astrogliogenesis, or astrocyte state conversion. Extending these novel roles of DOT1L in regulating astrocytes, it also impacts cortical astrocytes deriving from ventrally located, *Nkx2.1*-expressing progenitors. Most importantly, this study resolves diverse cell populations that feed into the astrocyte fate and state, and this plasticity argues in favour of a refined view that cortical astrocytes do not solely derive from APs after completion of neurogenesis.

## Material and Methods

### Sample collection and library preparation

Mouse pups were sacrificed by decapitation at embryonic day 12.5, 14.5 and 16.5 (E12.5, E14.5, E16.5) and postnatal day 0 (P0), their heads were transferred into ice-cold phosphate-buffered saline (PBS). The cerebral cortices were dissected and incubated in a papain and DNase containing solution for 30 minutes at 37°C. A single-cell suspension was obtained using the Papain Dissociation System (Worthington); all centrifugation steps were carried out at 4°C. The cell viability was confirmed to be higher than 90% before proceeding, cell concentration was measured using a Neubauer chamber. The cells were loaded into a Chromium X machine (10X Genomics) and the scRNA-seq library was prepared using Chromium Next GEM Single Cell 3⍰ Reagent Kits v3.1 (10X Genomics) following manufacturer’s instructions. The quality of the cDNA and libraries was assessed using a TapeStation 4150 (Agilent Technologies) and the concentration measured using a Qubit 2.0 (Thermo Fisher Scientific).

For the multiome (Nkx), pregnant mice carrying E16.5 embryos were sacrificed by cervical dislocation. Embryos were immediately placed in ice-cold phosphate-buffered saline (PBS) and decapitated, and collected heads were placed in ice-cold PBS. The region of interest was then dissected, nuclei were extracted, and libraries were prepared with Chromium X and the Chromium Next GEM Single Cell Multiome ATAC + Gene Expression kit (10x Genomics). The quality of the cDNA and libraries was assessed using a TapeStation 4150 (Agilent Technologies) and the concentration measured using a Qubit 2.0 (Thermo Fisher Scientific).

### Primary cultures of glial cells and EPZ5676 treatment

Primary cultures were obtained from postnatal day 3–4 (P3-P4) cerebral cortex of wild type C57BL/6 mice. Both hemispheres were dissected, meninges removed and mechanically dissociated using tweezers and by pipetting up and down with a decreasing micropipette tip diameter [29]. Centrifugation at 1200 rpm for 6 min at 4°C pelleted the cells, which were resuspended in Dulbecco’s Modified Eagle Medium (DMEM, Thermo Fisher Scientific, 42430-025) supplemented with 10% fetal bovine serum (Thermo Fisher Scientific, 10082147) and 100 μg/ml penicillin/streptomycin (Thermo Fisher Scientific, 15140122). The cells were plated on poly-lysine–coated (Sigma-Aldrich, P6407) T25 flasks and incubated at 37°C in a 5% CO_2_ atmosphere until confluence was reached. Medium change was conducted one day after plating and then after 3 days. To obtain an astrocyte-enriched culture (AEC), microglial depletion was obtained by 0.5 h of shaking under dark conditions at 180 rpm, and 6 h at 240 rpm to remove the oligodendrocyte precursor cells. Later, we rinsed the remaining confluent astrocyte layer twice with DPBS (Gibco 14190094), added 1,5 μl trypsin 0.25% (Thermo Fisher Scientific, 25200056) and plated on poly-lysine–coated glass coverslips [30]. Under every condition, the cells rested for at least 24 h before treatments, upon which AEC were treated either with 10 μM EPZ5676 (Active Biochemicals) or 1/1000 DMSO (Sigma-Aldrich, D8418) in complete medium for 3 or 7 days [31]. Medium change was conducted after 1.5 days.

### Immunofluorescent stainings and quantification

For *in vitro* analysis, AEC cells were fixed using 4% paraformaldehyde for 15 min at room temperature after 3 or 7 days DMSO or EPZ treatment. Before antibody incubation, fixed cells were permeabilized with 0.1% Triton-X in DPBS (Sigma-Aldrich, T8787) for 15 min, followed by 30 min of incubation with blocking solution (5% normal donkey serum in DPBS) (Abcam, ab7475). The cells were incubated overnight with specific primary antibodies diluted in blocking solution (SOX2 1:100, Santa Cruz, sc-365823; GFAP 1:500, Thermo Fisher Scientific, #13-0300). After removal of primary antibodies, the cells were washed three times with DPBS, followed by 4 h of incubation at room temperature with fluorescent secondary antibodies (Alexa 594, Alexa 488 (1:500, Thermo Fisher Scientific)). The cells were washed thrice in DPBS, incubated for 5 min with DAPI (1:1000, D9542, Sigma Aldrich) and washed thrice with PBS. Coverslips were mounted using Fluorescent Mounting Medium (SouthernBiotech, #0100-01).

For immunostainings on tissue sections, isolated P0 mouse brains for ctrl and Dot1l cKO were fixed with 4% paraformaldehyde (PFA 4%) for 24 h. The brains were incubated in 30% sucrose in DPBS overnight at 4°C and subsequently embedded in tissue freezing medium (Leica, #14020108926). Frozen embedded brains were cut in 14μm sections and mounted on SuperFrost Plus Microscope slides (Thermo Fisher Scientific). Dried slides were washed twice with DPBS, incubated 30 min with 0.25% Triton X in DPBS. Blocking was performed in 10% normal donkey serum, 0.1% Triton X in DPBS for 1 h, followed by the incubation with primary antibodies (ALDH1L1 1:1000, Abcam, ab277623) in blocking solution at 4°C overnight. After three washing steps in 0.1% Triton X in DPBS, incubation with secondary antibodies Alexa Fluor 594 (1:500, Thermo Fisher Scientific) in blocking solution for 1 h at RT, counterstaining with DAPI (1:1000, D9542, Sigma Aldrich), three additional washing steps with DPBS and mounting in Fluorescent Mounting Medium (SouthernBiotech, #0100-01) followed.

Images were taken using the Axioplan M2 fluorescent microscope (Zeiss) equipped with an Apotome.2 module or with Leica Sp8 confocal microscope. ZEN 3.0 and Fiji [32] were used for image processing. To analyze *in vitro* immunostaining, a two-way analysis of variance (ANOVA) was performed, followed by Sidak’s multiple comparisons test to evaluate differences between treatments (DMSO and EPZ) within each cellular phenotype. For *in vivo* histological analysis, we compared the number of positive cells between two conditions (Emx1-cre Dot1l cKO and WT), performing an unpaired Student’s *t*-test. In all cases, a sample size of N = 3 per group was used and a *p* value below 0.05 (*p* < 0.05) was considered as the parameter of significance. Statistical analysis and graph plotting were conducted using Prism GraphPad software (v6.07).

### Bioinformatics Analysis

#### scRNA preprocessing

Raw sequencing data was processed using the 10X Genomics Cell Ranger count pipeline (v7.1.0) with the mm10-2020-A transcriptome reference. Default pipeline settings were used to align reads and filter high-quality cells and, in the end, gene-by-cell count matrices were retrieved. Seurat (v5.2.0) [32] was further used for downstream quality control (QC). To remove low-quality cells, the mitochondrial percentage was calculated and cells with a high mitochondrial ratio (mito_ratio >= 0.12) were filtered out. Additionally, cells were filtered based on the other quality control criteria (UMI count per cell <= 1000, gene count < 500, and log10GenesPerUMI < 0.80). The datasets were normalised using SCTransform function followed by variable gene identification using FindVariableFeatures (selection.method = ‘vst’, nfeatures = 3000) function. Principal component analysis was performed using RunPCA function and *k*-nearest-neighbours graph was calculated for 40 components using FindNeighbours function. Cell clustering was performed using the standard Louvain algorithm for all resolutions from 0.1 – 0.8 to obtain high resolution clusters. For cluster annotation, marker gene identification was done using FindAllMarkers function with min.pct set to 0.25, logfc.threshold = 0.25 and test.use = Wilcoxon Rank Sum test. Manual cluster annotation was performed by integrating marker gene comparison against public databases (CellMarker2, UniProt, CellChat) with a custom curated set of brain-specific marker genes using our own script. Cell identities were then assigned by manual evaluation of the resulting marker gene signatures. To combine replicates, datasets were merged and combined using Seurat function JoinLayers.

For the multiome (Nkx), raw sequencing data was processed using 10X Genomics Cell Ranger ARC (v2.0.2) with the mm10 reference genome(mouse). The data was subsequently processed using the standard Seurat workflow. (R - v4.1.2, Seurat - v5.2.1).

### Cell type proportion analysis

The proportion differences between WT and Dot1l cKO cells per cluster were calculated using the R package scProportionTest (v0.0.0.9000). For each cluster, statistical significance was estimated by permutation testing with confidence intervals estimated by bootstrapping. Clusters exhibiting an observed fold difference of at least 0.40 or 0.58 and a false discovery rate (FDR) of <0.05 were considered to have statistically significant changes in cellular composition.

### Differential gene expression and GO term analysis

Differential gene expression analysis was performed separately for each cell type to identify genes that were significantly altered across genotypes (ctrl and Dot1l cKO). This was done by using Seurat FindMarkers function; genes with (log_2_ fold change (FC)| ≥ 0 or 0.25) and (adjusted *p*-value < 0.05) were considered significantly differentially expressed. Top 20 DEGs were shown using volcano plots generated by ggplot2 (v3.5.1 and R v4.4.3).

The resulting DEG lists were used for pathway enrichment analysis using clusterProfiler (v4.12.6) [33]. Enrichment was performed against Gene Ontology (GO) pathway databases (v3.19.1) for biological processes (BP), using an adjusted p-value threshold of < 0.05 to define significantly enriched pathways.

### Cell cycle analysis

Cell cycle phases for all datasets were identified using Seurat (v5.2.0) CellCycleScoring function. Cells were classified into G1, G2M and S phase. For a more detailed view of cell cycle phases ccAfv2 (v0.0.0.9000) R package was used [34]. This package classified cells into multiple phases (G0/G1, S, S/G2, G2M, M/Early G1) using PredictCellCycle function. Proportion of cells across each phase is shown using a bar plot.

### Cell type specific splice junction analysis

Aligned BAM files generated for each biological replicate using 10X Genomics Cell Ranger (v7.1.0) were used as input for downstream analyses. To facilitate batch processing, the default Cell Ranger BAM file names were renamed to sample-specific names. Cell type annotations and genotype information were extracted from a merged Seurat object processed in R (v4.4.3). Each cell barcode was then assigned a composite annotation (APs_cKO) generated by concatenating the cell type label with its genotype metadata. For each sample, a two-column tab-separated file was exported containing the cell barcode (stripped of the sample prefix) and its corresponding cell type label. These mapping files served as input for barcode-based read filtering in the subsequent step. Cell type-specific BAM files were generated using sinto filterbarcodes (sinto, v0.10.1). A bash script iterated over all barcode-to-cluster mapping files and identified the corresponding sample BAM file for each. For every sample, sinto was used to split the full BAM file into separate BAM files, one per cell type, based on the barcode annotations. Read extraction was parallelized using 20 CPU threads. Output BAM files were written into sample- and cell type-specific subdirectories.

To increase read depth and reduce replicate-specific noise, BAM files from biological replicates of the same timepoint, condition and cell type were merged using sambamba merge (sambamba, v1.0.1) with 20 threads. The pipeline first identified all unique timepoints and cell types present across the dataset. For each timepoint–cell type combination, if multiple replicate BAM files were found, they were merged into a single BAM file. In cases where only a single replicate existed, the BAM file was soft-linked to the output directory to maintain a consistent directory structure. All merged (or linked) BAM files were subsequently indexed using sambamba index (v1.0.1) to enable efficient random access in downstream steps.

To visualize RNA splice junction and read coverage across cell types at the *Dot1l* locus, sashimi plots were generated using rmats2sashimiplot (v3.0.0), a tool built on the MISO sashimi plotting framework. Cell type-specific BAM files, generated as described above, were used as input.

The genomic region of interest was defined on chromosome 10 (chr10:80,755,206– 80,768,801, + strand, mm10). Exon and intron scaling parameters were set to 1 and 5, respectively, to enhance intronic visualization. Junction read counts were filtered using a minimum count threshold of 1. Cell type grouping information was provided via a group configuration file, and custom colours were assigned per cell type and condition.

### Data integration using scVI

All WT and Dot1l cKO timepoints were combined individually and together using Scanpy (v1.9.5) concatenate function. Highly variable genes (HVGs) were identified using the default parameters of sc.pp.highly_variable_genes (flavour=‘cell_ranger’, n_top_genes=3000). The scVI (v0.20.3) model was then built using two hidden layers and a negative binomial gene likelihood and trained for a maximum of 400 epochs (default) with batch_key set to ‘day’ (timepoint) and labels_key=‘ct’ (cell type) using model_scvi.train. Finally, scVI-normalized latent representation was obtained using get_latent_representation, and the nearest neighbor graph was generated by sc.pp.neighbors (use_rep=‘X_scVI’) with default parameters. UMAP was constructed to visualize clusters using sp.tl.umap and later unsupervised Leiden clustering was used to identify clusters with multiple resolutions (0.5, 1, 2, 3, 5). No additional batch correction was performed beyond scVI integration.

### Trajectory analysis using CellRank2

Trajectory inference was performed using CellRank2 (v2.0.0) [40] on the integrated dataset (E12-P0) to model directed cell state transitions. Trajectories were inferred using the default parameters as described in the tutorial (https://cellrank.readthedocs.io/en/latest/notebooks/tutorials/index.html). A real-time kernel was constructed to capture transcriptional progression and infer cellular trajectories using RealTimeKernel.from_moscot function. In addition, RNA velocity-based kernels (VelocityKernel) and ConnectivityKernel were also computed and combined (combined_kernel = 0.8 * vk + 0.2 * ck) to assess the robustness of the inferred dynamics. CellRank’s built-in functions for identifying initial and terminal states were not used. Rather, root states were manually assigned to assess cell fate transitions from different starting points. Transition probabilities between cells were computed for each kernel, and the resulting lineage structure was compared across kernels.

### *In silico* gene perturbation using CellOracle

A machine learning framework CellOracle (v0.18.0 [41] with Python v3.8.20) was used to simulate gene perturbations. CellOracle used simulated gene expression changes to predict cellular transition trajectories across all transcription factors in both WT and Dot1l cKO conditions. Processed scRNA-seq data in anndata format with dimensionality reduction and clustering information was used as input across genotypes individually. Trajectories were recalculated using Pseudotime_calculator function. The key TFs identified earlier with DEGs (*Lhx2, Meis2, Nfia, Nfib, Tcf4, Sox6, Emx1, Creb5*) were used for *in silico* perturbation. In case of WT, TF expression was knocked down (expression = 0) while for Dot1l cKO condition TF expression was doubled. To calculate predicted states after perturbation oracle.simulate_shift function was used. oracle.calculate_p_mass function was used to estimate cell density across the embedding, it creates the grid around embedding and calculates how many cells fall near the grid. Positive or negative effect of perturbation post change is calculated by calculate_inner_product function, and the result of perturbation is shown on embedded grid using plot_inner_product_on_grid function.

## Results

### DOT1L limits gliogenesis and regulates astrocyte functions at synapses

Single-cell (sc)RNA-seq of postnatal (P0) Dot1l cKO (Dot1l^fl/fl^; Emx1^cre/+^) and WT (Dot1l^fl/fl^; Emx1^+/+^) forebrains identified two astrocyte fractions, Astrocytes I and II, clustering close to oligodendrocytes and apical progenitors (AP), which are considered as astrocyte precursors [35] (**Fig. 1a, S1a**). Both genotypes contributed to all observed cell clusters (**Fig. 1b**), however, cell proportions varied in several clusters (**Fig. 1c, d**). In accordance with other DOT1L loss-of-function (LOF) models *in vivo* and *in vitro* [25-28], showing increased neurogenesis, excitatory and inhibitory neuronal populations increased significantly, whereas neurogenic BPs and one out of two upper layer neuron clusters decreased (**Fig. 1d**). Notably, only the Astrocytes II population increased upon DOT1L LOF, whereas Astrocytes I did not change significantly in numbers. ALDH1L1 immunostaining at P0 (**Fig. 1e**) confirmed increased numbers of cortical astrocytes upon DOT1L LOF (**Fig. 1f**). Quantitative analyses of significantly differentially expressed genes (DEGs) revealed stronger effects of DOT1L LOF in Astrocyte I (33 increased, 147 decreased genes, 1.36 average LFC) compared to Astrocytes II (0 increased, 90 decreased genes, 1.33 average LFC) (**Fig. 1g**). Enrichment of functional GO-terms associated with the significant DEGs suggested that within Astrocytes I DOT1L LOF reduced cell proliferation, regulation of neurogenesis, dendrite and axon development. Gliogenesis as well as synaptic functions increased and decreased, respectively. Similarly, in Astrocytes II genes associated with dendrite/axon development and synaptic assembly/organisation decreased (**Fig. 1h**). The different responses in the two astrocyte populations upon DOT1L LOF could be caused by their origin from different progenitor lineages either expressing *Emx1* or *Olig2* [9]. Since we used *Emx1-cre* for the Dot1l cKO, an *Emx1*-positive or -negative origin could be a likely explanation. *Emx1* expression as proxy for *Emx1-cre* activity showed only slightly lower expression in Astrocytes II compared to Astrocytes I (**Fig. 1i**). However, *Emx1* expression levels were very low at P0 for both genotypes (**Fig. S1b**), which hampered discriminating Astrocytes I from II by a different origin (*Emx1*-positive, -negative) from a specific DOT1L LOF effect.

**Figure 1.**
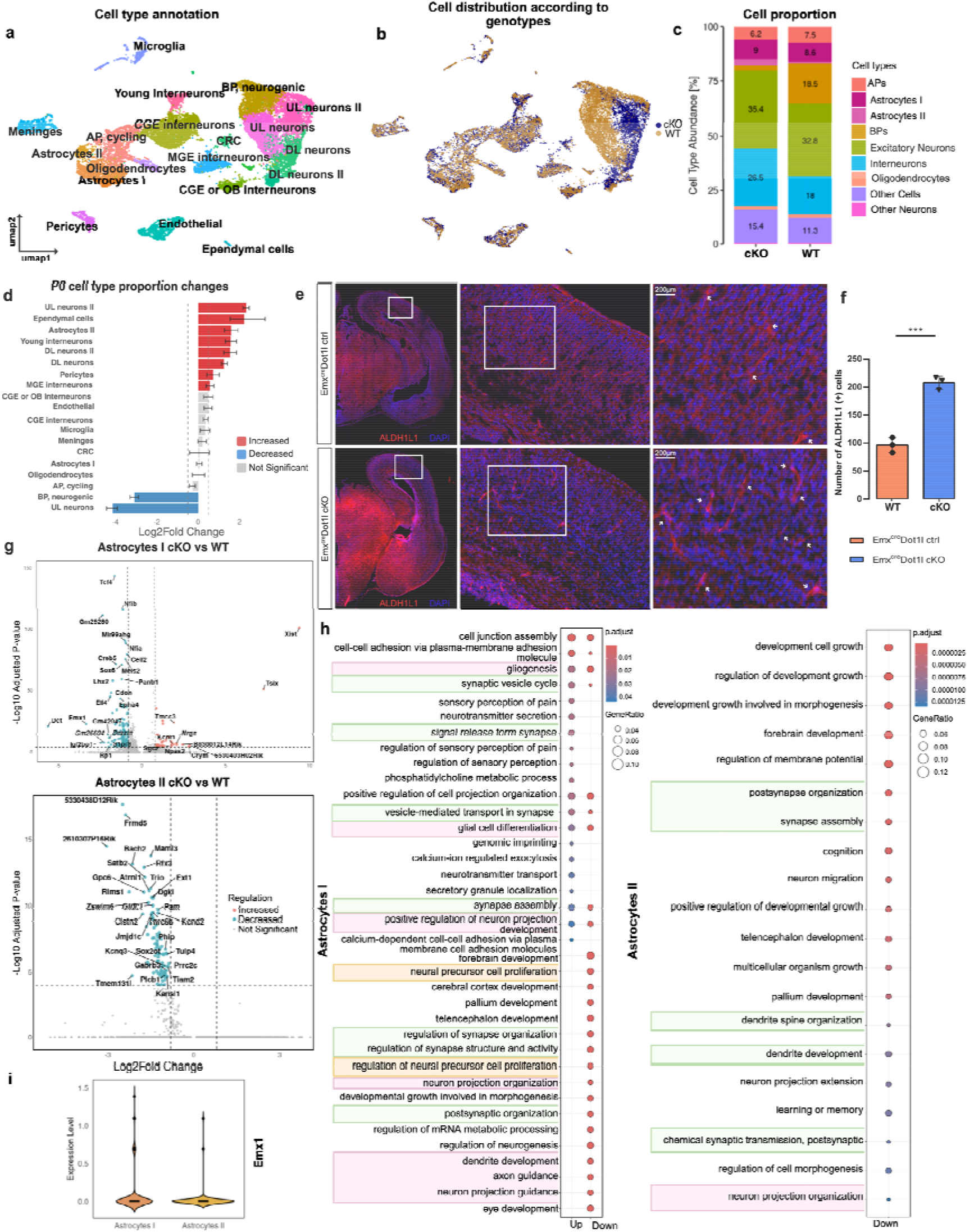
**a** UMAP representation of postal natal (P0) Emx cortex single cell transcriptomics dataset from *Emx1-cre* Dot1l cKO and WT control brains (n = 20,102 cells, WT n = 9,713, cKO n = 10,389). Colors represent different clusters. **b** UMAP representation of P0 cortex genotypes (WT and Dot1l cKO). **c** Bar plot showing the cell proportions across genotypes. **d** Bar plot illustrating cell proportion changes upon Dot1l cKO. Significantly increased or decreased cell types were defined with thresholds FDR < 0.05 and Log2Fold Change > 0.58. Error bars represent 95% confidence intervals from 1000 permutations. Red indicates significant increase, blue significant decrease, and grey no significant change. **e** Immunostaining of P0 cortex sections for ALDH1L1 in both ctrl and Dot1l cKO sections. Scale bars 200 μm. **f** Bar plot showing the quantification of ALDH1L1-postive cells in WT (n=3) and Dot1l cKO (n⍰=⍰3) samples with mean ± SEM. Individual data points represent biological replicates. Statistical significance was assessed by unpaired t-test. ***p < 0.001. **g** Volcano plot with genes that increased (pink) or decreased (blue) significantly upon the cKO for Astrocytes I and Astrocytes II. Horizontal dashed line indicates the significance threshold corresponding to an adjusted p-value of 1 × 10^−4^, and the vertical dashed lines show the Log2Fold Change threshold at 0.8 and −0.8. **h** GO biological process enrichment analysis of significant DEGs upon Dot1l cKO for Astrocytes I and Astrocytes II. Significantly enriched biological processes associated with different groups are indicated by color: green – synaptic processes, pink – axon-related and neurodevelopment processes and yellow - cell proliferation related processes. Enrichment was performed with Benjamini–Hochberg correction (*p* < 0.05, *q* < 0.2). **i** Violin plot illustrating the *Emx1* expression across both astrocyte populations. APs: Apical Progenitors, BPs: Basal Progenitors, cKO: conditional Knock Out

### DOT1L regulates numbers of astrocytes featuring translation, proliferation and homeostasis

To distill the differences between both two P0 astrocyte populations, we analysed the combined cell populations of WT and cKO cells in more detail. *Slc1a3, Tnc, Ptprz1, Slc1a2, Qk, Plpp3, Thyh1*, and *Ptn* showed relative specific expression in the Astrocytes I population, in comparison to Astrocytes II and to all other cell clusters of the P0 data set (**Fig. 2a**). In contrast to specific gene sets expressed in Astrocytes I, all top Astrocytes II markers were also expressed by Astrocytes I (**Fig. S1c**).

**Figure 2.**
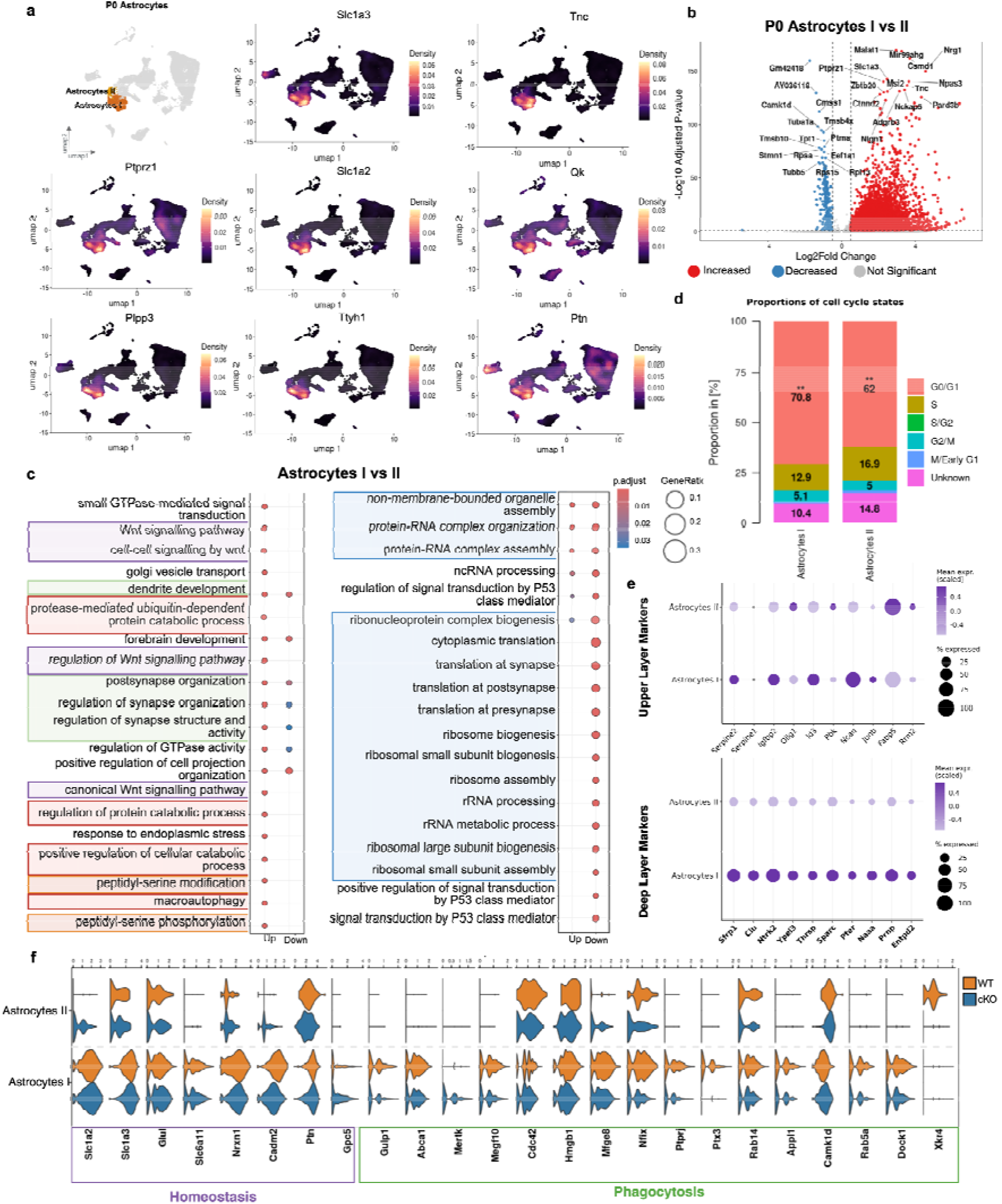
**a** Density plots representing marker gene expression enrichment in the P0 (WT and cKO) Astrocyte I population. **b** Volcano plot showing the significant differentially expressed genes between P0 Astrocytes I vs Astrocytes II (red: increased gene expression, blue: decreased gene expression, grey: not significant). Horizontal dashed line indicates the significance threshold of adjusted p-value = 0.05, and the vertical dashed lines show the Log2Fold Change threshold at 0.5 and −0.5. **c** GO enrichment analysis for biological processes of significant DEGs between P0 Astrocytes I vs Astrocytes II. Significantly enriched biological processes associated with different groups are indicated by colour: green – synaptic processes, purple – Wnt associated pathways, orange – peptidyl-serine related processes, red – protein degradation, and blue translation and biogenesis related processes. Enrichment was performed using Benjamini–Hochberg correction (*p* < 0.05, *q* < 0.2). **d** Bar plot showing the cell cycle states across genotypes represented in Astrocytes I and II. Statistical significance between Astrocytes I and II states was assessed using the Fischer test. Significance levels are denoted as ** (p < 0.01). **e** Dot plot elucidating the expression of upper layer (top) and deep layer (bottom) markers across P0 Astrocytes I and Astrocytes II. **f** Violin plot showing marker gene expression characteristic for phagocytosis and homeostasis for Astrocytes I and II across genotypes.

Significant DEGs between Astrocytes I and Astrocyte II mainly increased, in the data containing both genotypes combined (**Fig. 2b**), or split according to genotypes (**Fig. S2a, b**). In combined data, GO-term enrichment of significant DEGs indicated that Astrocytes I upregulated WNT- and GTPase signalling, serine modification and synaptic functions, whereas translation, protein-RNA complexes and p53-class signalling reduced in the combined data (**Fig. 2c**). To assess the DOT1L effects, GO terms were surveyed split according to genotypes (**Fig. S2c, d**). WNT- and GTPase signalling increased, whereas translation and p53-class signalling decreased in both genotypes, thus featuring characteristics of Astrocytes I compared to II. The alterations at the synapse and serine-modification associated with the DOT1L LOF, which also impaired activation of gliogenesis and regulation of neurogenesis in Astrocytes I.

Alterations of specific signalling pathways and translation were the most striking differences in gene expression programs between Astrocytes I and II, independent of DOT1L. WNT-signalling limits astrogenesis in the spinal cord [36]. We thus analysed the number of Astrocytes I and II in individual phases of the cell cycle, showing that Astrocytes I had a significantly higher number of cells in G0/G1 compared to II, in accordance with higher WNT signalling limiting Astrocytes I proliferation (**Fig. 2d**). WNT signalling also induces the reactive feature of astrocytes [37], but we did not reveal major differences in regard to a reactive phenotype (**Fig. S2e**). Astrocytes express distinct marker genes when localising to either deep or upper cortical layers [16]. The Astrocytes I expression program overlapped more with a deep layer as compared to an upper layer identity (**Fig. 2e**). Astrocytes are involved in phagocytosis [38,39], and Astrocytes I expressed more genes involved in this process compared to Astrocytes II (**Fig. 2f, S2f**), which seemed better described by homeostasis functions. Summarising the specific features, Astrocytes I are responsive to WNT signalling, restrict protein translation and proliferation, are capable of phagocytosis, and localise to deep layers of the cerebral cortex, whereas Astrocytes II engage in protein translation, proliferation and homoeostasis.

### Determination of marker genes for pre- and perinatal astrocyte subtypes

Our findings suggested functional differences between both P0 astrocyte populations, which might trace back to a heterogeneous origin, as recently suggested [9]. Alternatively, the differences between both populations might also indicate a state-switch between both populations. *Emx1, Olig2*, and *S100a11* discriminated different origins of astrocyte populations [9], but none of these were expressed in Astrocytes I or II (**Fig. S3a**). Similarly, Astrocytes I and II did not express markers for five proposed astrocyte subtypes present at P7 [9] (**Fig. S3b**). We concluded that P0 was neither suitable to align with marker gene expression for mature P7 astrocytes, nor for markers depicting early lineages. In search of developmental anchor points for Astrocytes I and II, we performed scRNAseq of WT and Dot1l cKO forebrains at E12.5 (**Fig. 3a**), E14.5 (**Fig. 3b**), and E16.5 (**Fig. 3c**). At E12.5 and E14.5 we did not detect distinct astrocyte clusters, as illustrated in the individual UMAP clustering. One cluster of Early Astrocytes, expressing canonical marker genes, appeared at E16.5 (**Fig. 3c, S4a**). However, these astrocytes did not express *Olig2, S100a11* and very few P7 astrocyte subtype markers (*Enkur, Igfbp5, Slc6a11, Slc7a10*, and *Id3*), similar to P0 (**Fig. S2b, S4b, c**), hampering classification of our astrocytes using these gene expression programs. We concluded that expression programs in astrocytes were heterogeneous and variable over the developmental time points, with postnatal gene signatures hardly represented in embryonic stages. We therefore determined embryonic and early postnatal astrocyte markers from published data sets, covering multiple developmental stages of the mouse brain, i.e. the embryonic stages E16.25, E16.5, E17.0, E17.5, and E18.0 (*LaManno*) [40], E17.0, E18.0, P1, and P4 (*DiBella*) [41], and E18.5 (*Tcf4-cKO*) [42]. We enriched these data with our E14.5, E16.5 and P0 data (*Emx*) as well as astrocytes identified in the cortex at E16.5 upon *Nkx2.1-cre* DOT1L LOF (*Nkx*). The marker genes identified from our P0 *Emx* cortex data set, together with the canonical markers, highlight astrocytes in prenatal stages better (**Fig. S5**) compared to those markers suggested from the other data sets [9,41,42]. As APs/RGCs share active genes with astrocytes, we included these cell types from our E14.5 and E16.5 data for discrimination. APs/RGCs signatures were weaker, but still active in astrocytes of all data sets (**Fig. S5**). Postnatal astrocyte markers [9], were weakly indicative of embryonic astrocytes, with only subsets of genes from each subgroup being active. The similarity of embryonal and early postnatal astrocytic transcription programs was evident by comparing the E16.5 to P0 astrocyte marker genes, in which E16.5 Early Astrocytes had similar gene expression programs compared to P0. Together this analysis indicated different expression profiles in embryonic and early postnatal compared to postnatal astrocytes, and confirmed Early Astrocytes at E16.5 as astrocytes, and not APs/RGCs.

**Figure 3.**
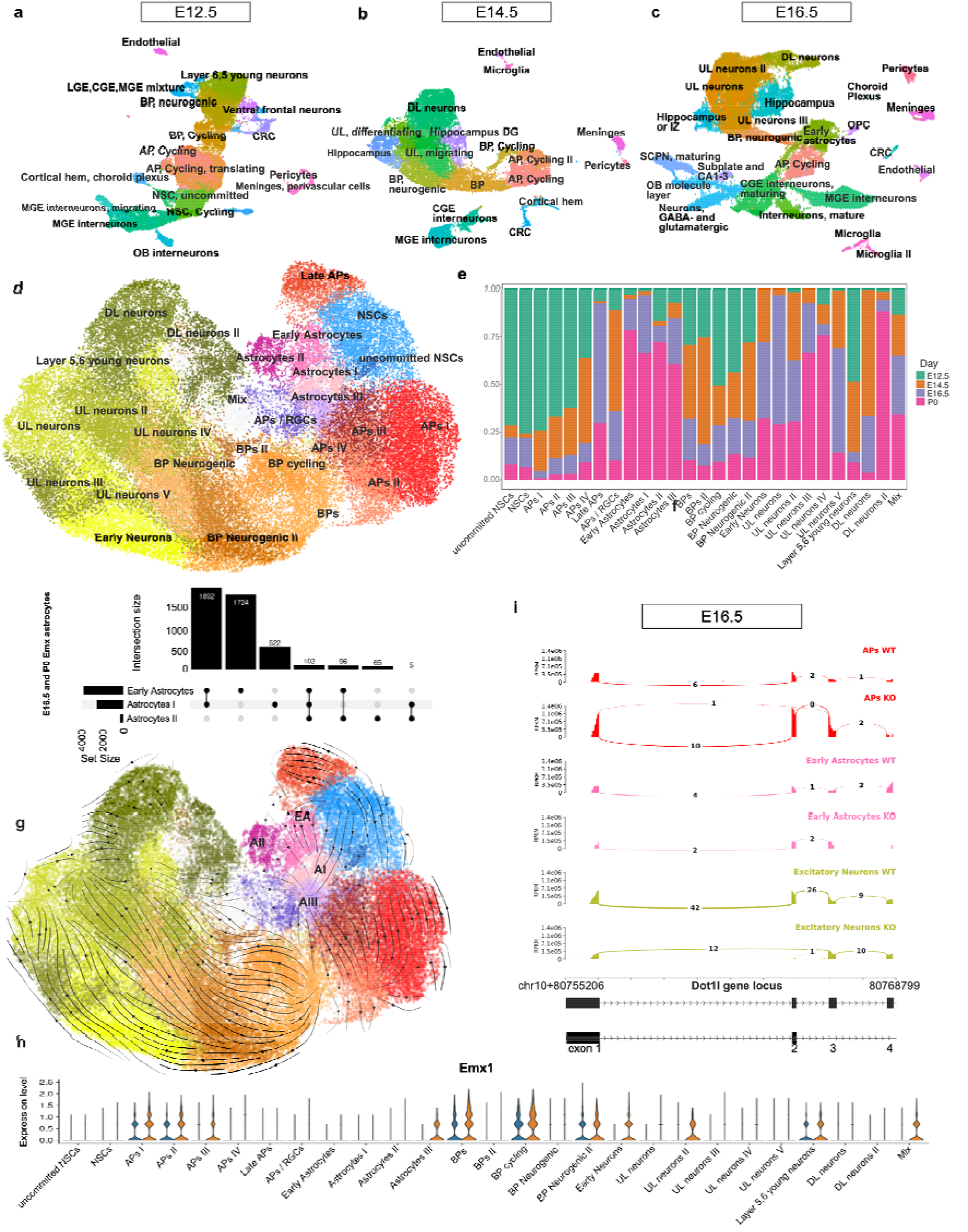
**a-c** UMAP representation of single cell transcriptomics data at E12.5 **(a)**, E14.5 **(b)**, and E16.5 **(c)** from *Emx1-cre* Dot1l cKO and control brains. **d** UMAP embedding of integrated single cell transcriptomics dataset (n=63,904) showing dorsal telencephalon derived cell types across 4 time points (E12.5, E14.5, E16.5, and P0). UMAP coloured by cell types. **e** Bar plot illustrating the proportion of cells corresponding to different time points in each cell type. **f** Upset plot showing the overlap of significant DEGs between *Emx1-cre* Dot1l cKO E16.5 (Early Astrocytes) and P0 Astrocytes I and II. **g** Cell differentiation is projected using streamlines based on RealTimeKernel on top of the UMAP embedding. Arrows suggest the direction of cell differentiation. Cluster labels for astrocytes are shown on top of the UMAP, abbreviations were used for readability (EA – Early Astrocytes, AI – Astrocytes I, AII – Astrocytes II, AIII – Astrocytes III). ***h*** *Emx1* expression across all cell types of integrated dataset (E12.5-P0) across genotypes (WT – orange, cKO – blue). **i** Sashimi plots displaying the RNA-seq read coverage (RPKM) and splice junction usage at the *Dot1l* gene locus (chr10: 80,755,206–80,768,799) in different cell types for both WT and cKO conditions at E16.5. Filled areas represent per-base read depth scaled in RPKM (0 – 1.4×10^6^). Arcs connecting exonic regions represent split reads spanning splice junctions, with the number above each arc indicating the number of junction-supporting reads. The gene model at the bottom depicts annotated *Dot1l*, with black boxes representing exons and horizontal arrows indicating intron direction on the forward (+) strand. Exons 1–4 are labelled.

### Emx1-cre expression discriminates dual origin of astrocytes

We aimed to reproduce dual lineage origin for astrocytes as recently reported [9] in our data by assessing lineage trajectories of astrocytes. For this we integrated our scRNA-seq data from E12.5, E14.5, E16.5, and P0 for both genotypes into one dataset (**Fig. 3d**), excluding cell types that derive from progenitors residing mainly outside the cortical plate (interneurons, microglia, pericytes, endothelial cells, etc.). Canonical marker gene expression confirmed the presence of different clusters of astrocytes (**Fig. S6a**). As a result of the data integration, several cell clusters appeared in a refined organisation, with contribution of earlier stages, i.e. E12.5 and E14.5 to the astrocyte population. Accordingly, we identified RGCs/APs, Early Astrocytes, and Astrocytes I, II, and III, all of which had contribution from cells from all four time points (**Fig. 3e**). Except for the RGCs/APs, P0 cells contributed the most cells to the respective astrocyte clusters. This integrated data set allowed us to link maturing astrocytes at P0 to respective progenitors.

In regard to subtype specification, Astrocytes II in the integrated data kept their distinct homeostatic features, whereas phagocytosis features were present in Astrocytes I and Astrocytes III (**Fig. S6b**). Unbiased functional GO-term analysis showed distinct processes active in the respective astrocyte populations, exposing differences in translation, proliferation, metabolism, mRNA processing and synaptic organisation. In regard to upregulated processes, Early Astrocytes overlapped more with Astrocytes I compared to II (**Fig. S6c**), similar to the number of shared expressed genes, which was higher with Astrocytes I compared to II (**Fig. 3f**). In terms of upregulated processes, Astrocytes III only had two processes as unique identifiers, both of which related to mitochondrial function. This astrocyte type had the larger overlap with Early Astrocytes and Astrocytes I, compared to Astrocyte II.

For resolving the cell lineage trajectory of astrocytes, we used the Cellrank algorithm [43] on the integrated data set with both genotypes, which predicted an astrocyte lineage from NSCs to Early Astrocytes, Astrocytes I and Astrocytes III, in addition to the established origin of astrocytes from APs, which mainly connected to Astrocytes III (**Fig. 3g**). Late APs, mainly coming from the E16.5 data set (**Fig. 3e**), also derived from NSCs and seemed to feed specifically into the Astrocyte II population. As NSCs as well as Late APs did not express *Emx1* (**Fig. 3h**), in contrast to APs I-III in the integrated data (E12-P0), we also suspected traces of a dual astrocyte origin in our data. Since *Emx1-cre* activity should result in the absence of the floxed exon 2 of *Dot1l* in our cKOs, we used Sashimi plots revealing presence/absence of specific splice junctions. At E12.5, cells from the excitatory lineage retained the exon 1-2 junctions only in the WT condition, reflecting *Emx1* activity and DOT1L LOF within the cKO animals (**Fig. S7a**). NSCs at this stage did not express *Emx1-cre*, whereas APs did. At E16.5, *Dot1l* exon 1-2 junctions were nearly absent in excitatory neurons, while Early Astrocytes kept the junction in both genotypes. Similarly, APs in the cKO and WT samples retained the exon 1-2 junctions (**Fig. 3i**). This suggested an *Emx1*-negative origin of Early Astrocytes at E16.5, either from NSCs or Late APs, which did not activate *Emx1* expression compared to E12.5 APs (**Fig. S7a**). Importantly, this finding supports a dual origin of astrocytes from *Emx1*-positive, i.e. APs, and *Emx1*-negative, i.e. NSCs and Late APs, cortical precursors, in accordance with other’s data [9]. Our data also show that *Emx1*-negative NSCs refill the AP pool at mid/late neurogenesis. We concluded that earlier views of astrocyte’s origin from APs that switch from neuro- to astrogliogenesis needs refinement in a way that considers different progenitor populations, NSCs and APs, contributing to the astrocyte populations. The lineage trajectory suggested that Astrocytes I had mixed origins from NSCs and APs, whereas Astrocytes II were most likely connected to Late APs. This is in accordance with the larger set of significant DEGs upon Dot1l cKO in Astrocytes I, deriving from APs of the *Emx1*-positive lineage, and Astrocytes II probably deriving from the *Emx1*-negative lineage. However, it left the question open, why Astrocytes II had increased cell numbers and a set of DEGs despite seemingly not being directly affected by the cKO.

### DOT1L controls TF expression in embryonic and early postnatal astrocytes

As the trajectory indicated that Early Astrocytes originated from an *Emx1*-negative, and thus DOT1L-retaining cell lineage, we explored the impact of DOT1L LOF at E16.5, containing this astrocyte population. Upon DOT1L LOF, the Early Astrocytes at E16.5 did not change significantly in numbers (**Fig. 4a, b**), but had a set of DEGs (**Fig. 4c**). Among the genes with significantly decreased expression, we identified *Tcf4, Lhx2, Emx1* and *Igfbp2*, all of which also decreased at P0 upon DOT1L LOF within the Astrocytes I (**Fig. 1g, 4c**). Comparison of the significant DEGs upon DOT1L LOF at E16.5 and P0 in the represented astrocyte populations revealed a higher fraction of DEGs shared between Early Astrocytes and Astrocytes I, whereas the shared fraction between Early Astrocytes and Astrocytes II was smaller (**Fig. 4d**). Consistently, Early Astrocytes also expressed marker genes for homeostasis and phagocytosis, supporting their similarity to the Astrocytes I population (**Fig. 4e**). The transcriptional relationship between Early Astrocytes and Astrocytes I support the view that *Emx1*-negative NSCs generate an early generation of astrocytes that feed the Astrocytes I population observed at P0. But this Astrocytes I pool is also fed by APs, according to the classical view. However, if Early Astrocytes would have solely derived from *Emx1*-negative NSCs, it remained unexplained why they had DEGs, among which we find “classical” DOT1L-targets, which are also regulated in other CNS-residing cells (e.g. *Lhx2, Tcf4*). This suggested that *Emx1*-lineage APs contributed not only to Astrocytes I, but also to Early Astrocytes. Alternatively, we speculated that the identified astrocyte populations could change their identities, eventually converging on a population of mixed origin.

**Figure 4.**
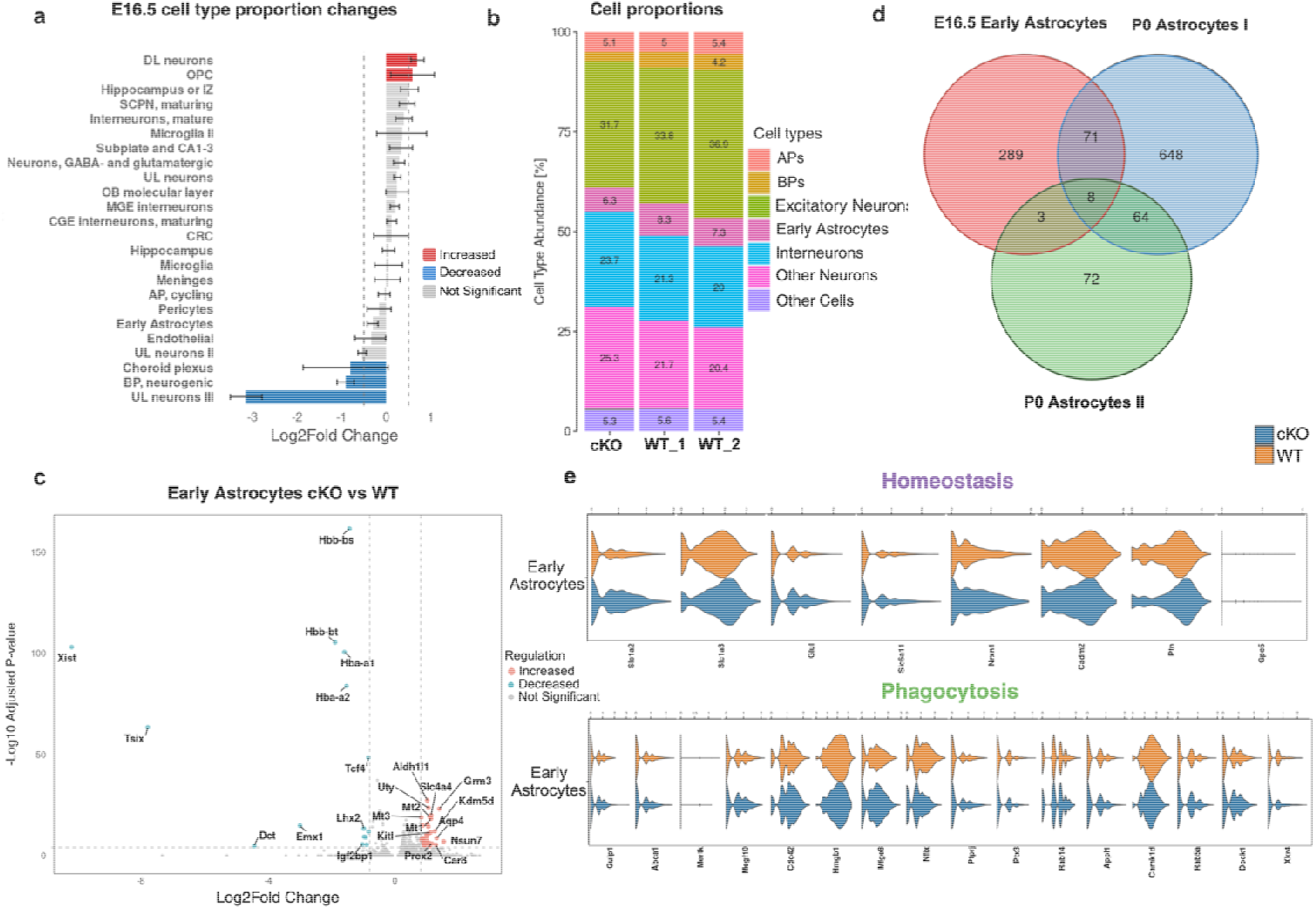
**a** Bar plot illustrating the cell proportions changes upon Dot1l cKO in E16.5. Significantly increased or decreased cell types were defined with thresholds FDR < 0.05 and Log2Fold change > 0.58. Error bars represent the 95% confidence intervals, calculated from 1000 permutations. Red indicates significant increase, blue significant decrease, and grey no significant change. **b** Fractions of cell type proportions across genotypes. **c** Volcano plot with genes that increased (pink) or decreased (blue) significantly upon Dot1l cKO in Early Astrocytes. Horizontal dashed line indicates the significance threshold corresponding to an adjusted p-value of 1 × 10^−4^, and the vertical dashed line shows the Log2Fold Change threshold at 0.8 and −0.8. **d** Venn diagram showing the overlap of DEGs across E16.5 and P0 astrocyte populations upon Dot1l cKO. **e** Violin plots displaying the expression of marker genes for phagocytosis and homeostasis across genotypes in Early Astrocytes at E16.5.

### DOT1L regulates astrogenesis and astrocyte state conversion and limits re-activation of NSC identity

Our data supported heterogenic origins of astrocytes from variable stem cell pools, i.e. *Emx1*-negative and -positive stem cells, NSCs and APs. However, the cell trajectories (**Fig. 3g**) seemingly lacked resolution to identify potential state conversions between progenitors and astrocytes, as well as within the astrocyte populations, explaining differences in the response to DOT1L LOF. To tackle this question, we analysed cell fates with a higher resolution. Using alluvial plots, we predicted and resolved the cell transition potential between stem cells and astrocytes. Centred view points on NSCs and APs indicated that all WT stem cells contributed to all astrocyte populations, as well as to the progenitor pools themselves, to variable extent and with different dynamics during development from E12.5 till P0 (**Fig. 5a**). If NSCs were largely *Emx1*-negative at E12.5 (**Fig. 3h**), they provided a fraction of *Emx1*-negative astrocytes. APs as *Emx1*-positive progenitors, also contributed and built up the *Emx1*-positive fraction of astrocytes. Notably, NSCs contributed to APs and vice versa, suggesting that also the stem cell pool consisted of cells from both lineages, at least in later stages of embryonic development.

**Figure 5.**
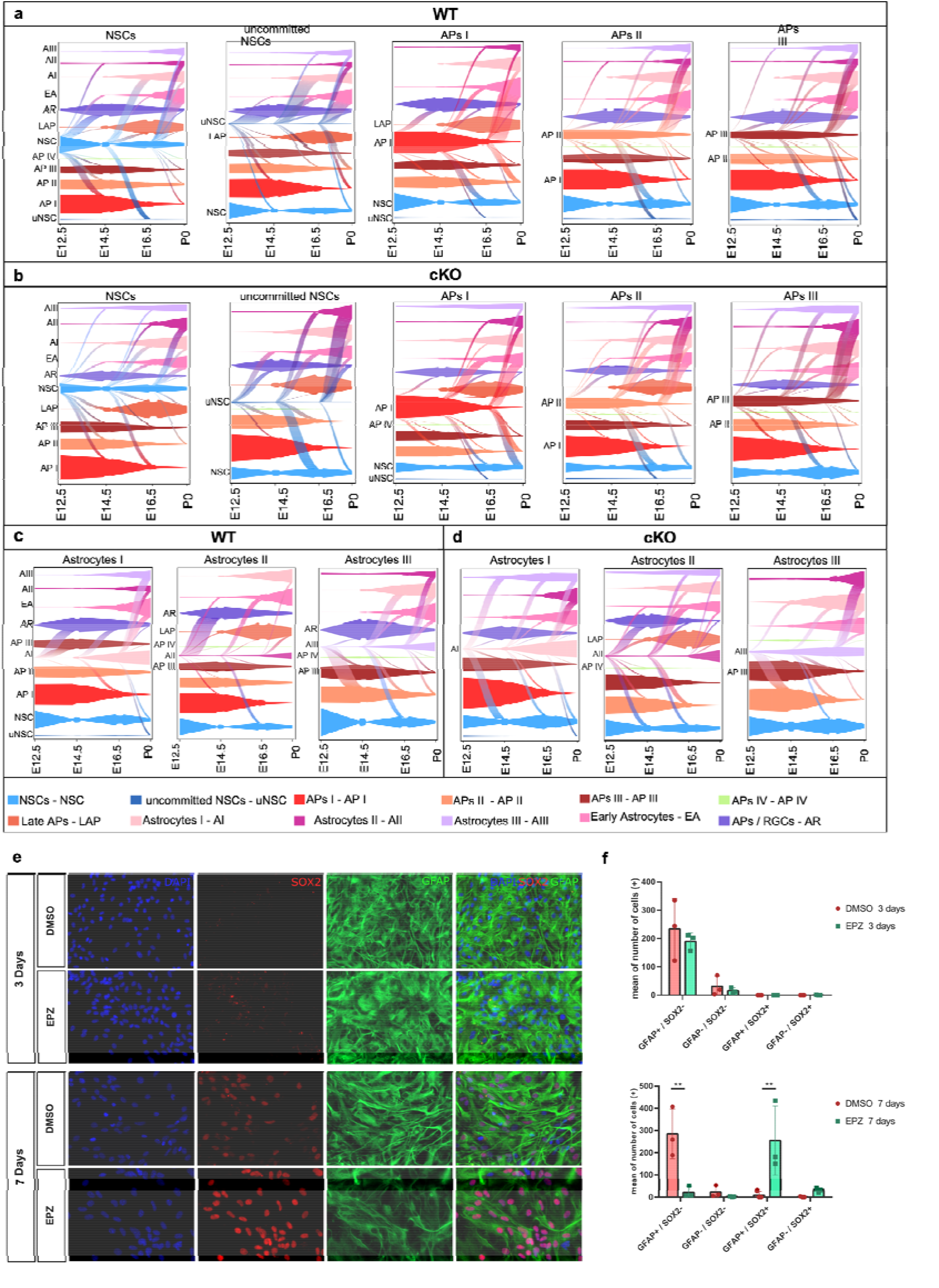
**a** Alluvial plot showing the transition probabilities across different WT cell groups (NSCs, uncommitted NSCs, APs I, APs II, APs III), and **b** across cKO cells. **c** Alluvial plot illustrating the transition probabilities across WT Astrocytes I-III, and **d** across cKO Astrocytes I-III. Colours represent different clusters and flows connecting cell types describe lineage transitions, with the flow width indicating predicted probability. All the clusters for which the cell fate transition is shown are labelled on the Y axis. Timeline of the transition is shown with time points present on the X axis. Abbreviations of cell labels were used for readability (NSC – NSCs, uNSC – uncommitted NSCs, AP I – APs I, AP II – APs II, AP III – APs III, AP IV – APs IV, LAP – Late APs, Ast P – Astrocyte Progenitors, EA – Early Astrocytes, AI – Astrocytes I, AII – Astrocytes II, AIII – Astrocytes III). **e** Immunostaining with SOX2 and GFAP, detecting astrocytes expressing stem cell marker SOX2 at 3 and 7 days after DOT1L inhibition or DMSO control. **f** Quantification of GFAP-negative/SOX2-negative, SOX2-/GFAP-negative, SOX2−/GFAP-positive, GFAP-negative/SOX2-positive cells after 3 days (top), and 7 days (bottom) of DOT1L inhibition or DMSO treatment. Control (n⍰=⍰3) and DOT1L inhibition (n⍰=⍰3) as mean ± SEM. Significance was estimated using two-way ANOVA.

The trajectory predictions not only confirmed for the WT a high probability of both, NSCs and APs differentiating towards astrocytes (**Fig. 5a**). Unprecedented, the analysis also showed that NSCs contributed more to the astrocyte lineage in the early developmental stages E12.5 and E14.5, compared to APs. At later development stages, E16.5 onwards, our data predicted the opposite, a stronger contribution of APs than NSCs to astrocytes, in accordance with the prevailing view of APs switching to astrogliogenesis.

If the dynamics in cell state conversion/differentiation would be altered upon DOT1L LOF, the observed differences in the repertoire of affected genes or cell numbers (**Fig. 1d, g**) would be explainable. Indeed, *Dot1l* cKO altered the probabilities of stem cells differentiation towards astrocytes (**Fig. 5b**). Both NSC populations contributed less to Astrocytes I, but much more to the Astrocyte II state. Specifically, at E16.5, NSCs contributed less to Early Astrocytes in the cKO than in the WT. APs showed subtype specific differences, with APs I keeping grossly the WT fates, but APs II contributing more to Astrocytes I, and APs III more to Astrocytes II. Summarising, DOT1L LOF altered the probabilities of cell trajectories from NSC and APs towards different astrocyte populations, strongly favouring Astrocytes II fates, which was in support of the observed quantitative differences in this population (**Fig. 1d**). As multiple stem cell types contributed to variable extends to all astrocyte populations, DOT1L LOF could result in the observed variable features, including the differences in cell numbers and DEGs.

The observed cell state plasticity prompted us to predict the probability of the mature astrocyte populations I-III for changing their state or fate. From the astrocytes’ perspective (**Fig. 5c**), several conversions between astrocyte states in the WT appeared likely, including activation of progenitor programs as well as of other maturation signatures. This finding indicated astrocytes’ state plasticity during development. Again, DOT1L LOF impacted this plasticity. Astrocytes I had less potential to re-activate expression programs characteristic of Early Astrocytes compared to the WT. But they convert with higher probability to Astrocyte II, and are predicted to activate Astrocyte III identity earlier compared to WT (**Fig. 5d**). Similarly, Astrocyte II had increased potential to convert to Early Astrocytes and Astrocytes III in cKO compared to WT condition. Astrocytes III most prominently activated Astrocyte II expression programs upon DOT1L LOF. We concluded that increased cell numbers of Astrocytes II upon Dot1l cKO was also attributable to state conversion from mature astrocyte populations, in addition to increased generation from specific NSCs and APs.

Interestingly, we also observed a proportion of astrocytes that was predicted to de-differentiate to stem cell states, including RGCs/APs, APs and NSCs, and that loss of DOT1L resulted in reactivation of stem cell states (**Fig. 5c, d**). DOT1L LOF increased de-differentiation from Astrocytes I towards NSCs compared to WT, and Astrocytes II had increased potential towards AP II and NSCs earlier in development, whereas Astrocytes III had less probability to convert towards NSCs (**Fig. 5d**). In support of DOT1L’s role in limiting conversion from NSCs to astrocytes, pharmacological DOT1L inhibition of primary astrocytes *in vitro* with EPZ5676/pinomenostat increased the fraction of SOX2-expressing GFAP-positive cells, while decreasing the number of GFAP-positive cells not expressing the stem cell marker SOX2 compared to vehicle-treated astrocyte cultures (**Fig. 5e, f**).

In summary, DOT1L functions in astrogliogenesis by preventing premature appearance of astrocytes with limited phagocytosis activity, and by timing astrocyte differentiation along a refined developmental trajectory including NSCs in addition to APs. DOT1L limits expression of mature astrocyte gene programs as well as de-differentiation towards stem cell states.

### In silico modelling reveals DOT1L-targeted TFs affecting astrocytes fate and state

To expose the underlying mechanisms by which DOT1L LOF alters astrogliogenesis and astrocyte maturation, we explored TFs affected in Dot1l cKOs, assuming them as main driving forces. A set of TFs, including *Tcf4, Lhx2, Emx1, Nfia, Nfib, Meis2, Sox6*, and *Creb5*, decreased upon Dot1l cKO at P0, and at E16.5 (**Fig. 6a**). Notably, while these TF were significant in Early Astrocytes and Astrocytes I, they were DEGs in Astrocytes II, but not within the significant fraction. This might reflect again the heterogeneous origin of Astrocytes II from *Emx1*-positive/DOT1L-negative as well as *Emx1*-negative/DOT1L-positive progenitors. To test the contribution of these plausible candidate TF to the observed phenotypes, we used *in silico* gene perturbation to assess the effects of either deleting these TF from our WT data, or by adding these to our cKO data, using CellOracle [44].

**Figure 6.**
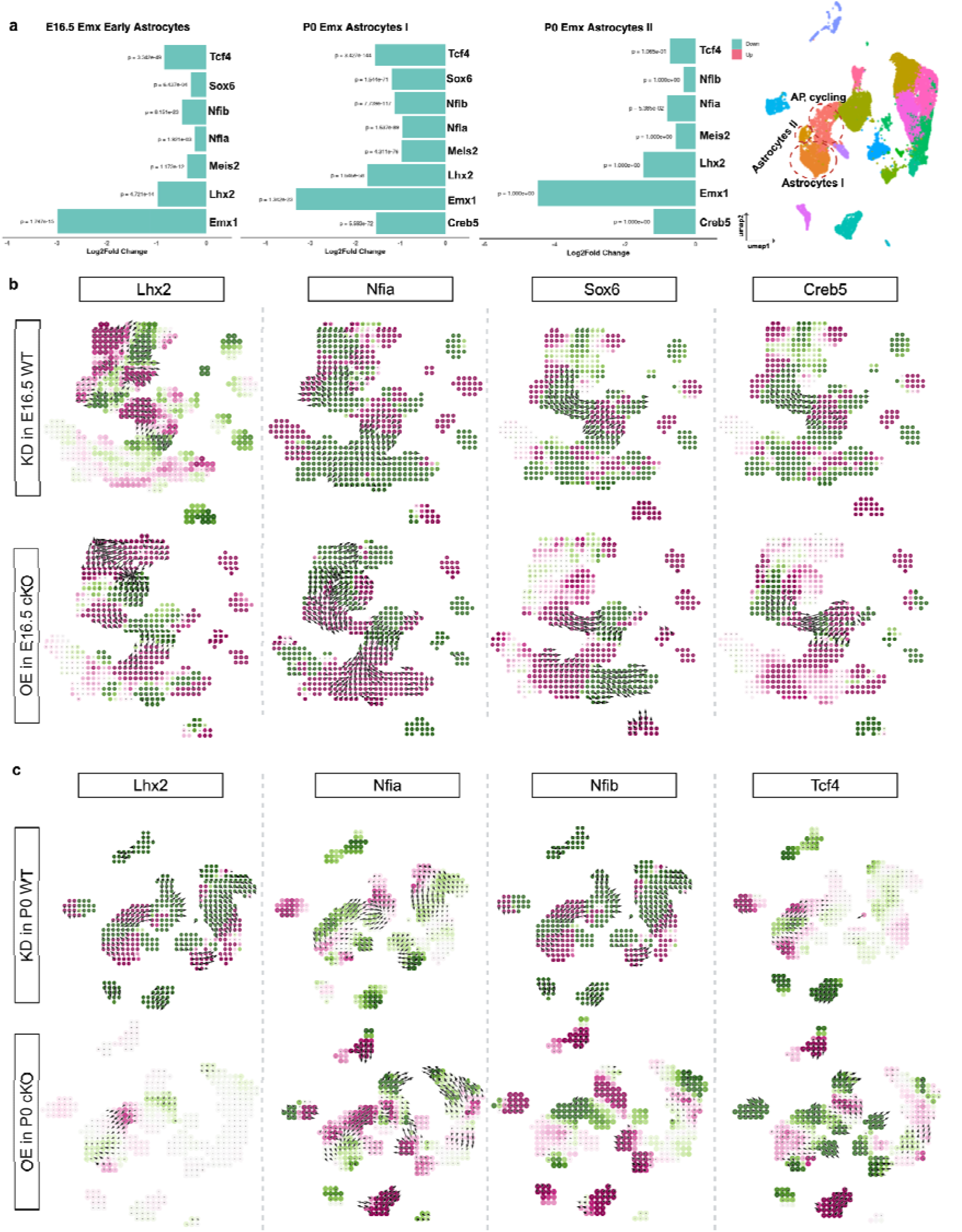
**a** TFs changing upon Dot1l cKO in Early Astrocytes (E16.5), and Astrocytes I and II (P0). Change is shown using Log2Fold Change and P-value significance is shown on the bars.Green color denotes decrease upon the Dot1l cKO. **b** Simulation vectors show the predicted transition in cell states after *Lhx2, Nfia, Sox6*, and *Creb5* knockdown (KD) (top), for each gene individually in E16.5 WT dataset, and their OE (bottom) in E16.5 cKO dataset representing the altered likelihood of a cell state change post perturbation. **c** Simulation vectors show the predicted transition in cell states after *Nfia, Nfib*, and *Tcf4* KD (top) individually in P0 WT dataset, and OE (bottom) in P0 cKO dataset representing the altered likelihood of a cell state change post perturbation. Pink and green denote the effects of perturbation on lineage trajectories. Pink indicates a deviation from the expected developmental trajectory, whereas green indicates that the differentiation trajectory remains aligned with the expected developmental lineage. The arrows represent the predicted shift in cell identity in response to the TF perturbation. Cluster localisation as indicated in the UMAP top right corner. KD: knockdown, OE: overexpression

We assessed perturbations of TF expression at E16.5, deducing whether APs produced more or less Early Astrocytes, and at P0, analysing Astrocytes I to II conversion. The perturbations were reciprocal, i.e. reduced expression of TF in WT cells, and increased in cKO cells, respectively (**Fig. 6b, c, Table S1**). We tested at E16.5, whether DOT1L promoted astrogenesis (i.e. AP to Early Astrocyte differentiation) by activating *Lhx2, Nfia, Sox6*, and *Creb5* expression, as these were reduced upon DOT1L LOF. Indeed, their reduction in WT cells *in silico* was predicted to favour the AP over Early Astrocyte state, and their overexpression in cKO cells favoured Early Astrocyte differentiation from APs (**Fig. 6b, Table S1**). At P0, reduction of *Lhx2, Nfib*, and *Tcf4* in WT favoured the transition of APs to Astrocytes II, and the increase in cKO cells predicted re-activation of the AP state for *Nfib* and *Tcf4* (**Fig. 6c, Table S1**). In regard to the conversion of the Astrocytes I and II states, *Lhx2, Nfia, Nfib* and *Tcf4* reduction in WT cells also promoted this transition, and increased levels of *Nfia, Nfib* and *Tcf4* in cKO cells favoured the Astrocytes I state. TCF4 might only play a minor role in astrogliogenesis [45]. Since it occurred prominently in our data, we used a public scRNA-seq data set to test its role in this process. Applying the same CellOracle analysis of E18.5 scRNA-seq data from *Tcf4* LOF mice [42], reproduced our observation in WT cells that *Nfib, Lhx2, Meis2* and *Tcf4* favoured astrocyte differentiation via APs (**Fig. S8**). However, there was a limited effect on astrogenesis upon overexpression of the set of TF in cKO cells, in accordance with the restricted impact of *Tcf4* cKO on astrogliogenesis. Nonetheless, we observed that astrocytes converted from one cluster to the other upon overexpression of *Meis2*, and *Tcf4* in cKO cells (**Fig. S8**). Together, *in silico* modelling supported a role of DOT1L in regulating astrogliogenesis by controlling the expression of specific set of transcription factors, including *Lhx2, Nfia, Sox6*, and *Creb5*, that partly impact neurogenesis as well [46-49]. Regulation of *Nfib, Sox6, Meis2*, and *Tcf4* by DOT1L is implicated in preserving mature astrocyte states.

### DOT1L stabilises S-phase expression programs at E12. 5

In support of our refined view that astrocytes derive not only from APs switching from neuro-to astrogliogenesis, we noticed equal numbers of Early Astrocytes at E16.5 and Astrocytes I at P0, despite decreased numbers of neurogenic BPs and upper layer neurons (**Fig. 1d, 4a**). The latter are indicative of altered late neurogenesis. Cell proportions at the respective time points in our scRNA data showed a reduction of NSCs, and cycling APs at E12.5, but increased numbers of differentiating APs and BPs. Indicative for premature differentiation, we identified fewer BPs at E14.5 upon DOT1L LOF (**Fig. 7a**). At E16.5, APs were not different in numbers between the two genotypes, which might point towards a potential for recovery or refill of APs (**Fig. 4a**). As DOT1L controls cell cycle progression [27,50], we investigated differences in the proliferation rates across genotypes, as increased cell proliferation would be a prerequisite to compensate for reduced numbers of stem cells. Cell cycle phases in the integrated data set (E12.5-P0) in the individual cell populations across genotypes (**Fig. 7b**) indicated that NSCs, AP III and late APs had increased numbers of cells in S- or G2/M-phases upon DOT1L LOF. Thus, a refill mechanism to reconstitute potentially reduced numbers of *Emx1*-positive/DOT1L-negative progenitors in the AP lineage might be possible. In addition to the progenitor populations, Early Astrocytes and Astrocytes II seemed less proliferative upon DOT1L LOF compared to controls, whereas the other astrocyte populations were mainly unaffected.

**Figure 7.**
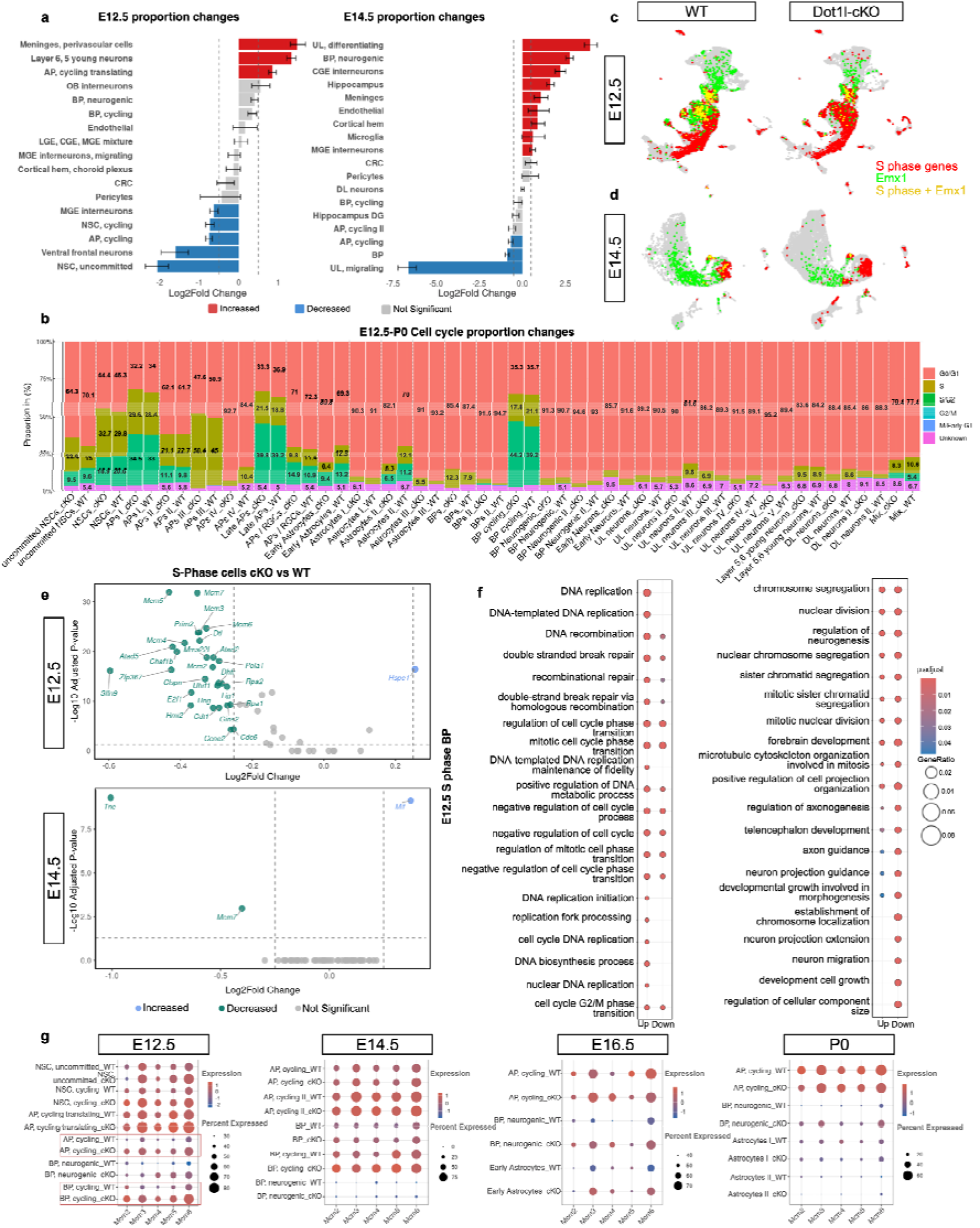
**a** Bar plot illustrating cell proportion changes upon Dot1l cKO at E12.5 (left) and E14.5 (right). Significantly increased or decreased cell types were defined with thresholds FDR < 0.05 and Log2Fold Change > 0.58. Error bars represent 95% confidence intervals from 1000 permutations. Red indicates significant increase, blue significant decrease, and grey no significant change. **b** Bar plot showing the cell cycle proportions (G0/G1, S, G2/M, M/Early G1, Unknown) across genotypes (WT n = 34,755, cKO n = 29,149) in the integrated dataset (E12.5-P0). **c** Feature plot illustrating the coexpression of S-phase genes and *Emx1* across genotypes in E12.5 dataset, and **d** in E14.5 dataset. Red, green and yellow colours show the expression of S-phase genes, *Emx1*, and S-phase+/*Emx1*+, respectively. **e** Volcano plot depicting the increased (blue), decreased (green) and not significant altered S-phase genes (grey) for E12.5 (top), and E14.5 (bottom). Horizontal dashed line indicates the significance threshold of adjusted p-value = 0.05, and the vertical dashed lines show the Log2Fold Change threshold at 0.25 and −0.25. **f** GO term enrichment analysis for biological process of E12.5 S-phase genes. Enrichment was performed with Benjamini–Hochberg correction (*p* < 0.05, *q* < 0.2). **g** Dot plot showing the expression of Mcm family members (*Mcm2-6*) in different cell types across different time points. Cell types for each time point are present on the Y axis and gene names are present on the X axis. The dot size represents the percentage of cells having the expression, and the colour indicates the scaled average expression calculated from the samples.

Extending our analysis of DOT1L’s potential impact on cycling cells, we identified in our scRNA data all cells in S-phase (E12.5, E14.5, E16.5, P0, separated) that presented as *Emx1*-positive and -negative (**Fig. 7c, d, S9a**). In S-phase cells, DOT1L LOF increased gene expression in the cKO, mostly at E12.5, very mildly at E14.5, and without significant changes at E16.5 or P0 (**Fig. 7e, S9b**). GO-term enrichment analysis indicated the activation of DNA repair processes, and G2/M progression checkpoints upon DOT1L LOF at E12.5 (**Fig. 7f**), hallmarks of replication stress, which could result in increased apoptotic elimination of the affected cells. On the other hand, cycling cells at E12.5 increased expression of MCM family members, involved in DNA damage repair (**Fig. 7g**). These data are in accordance with the hypothesis that stem cells, unaffected by the cKO, could partially compensate for early reduction in the progenitor pool affected by the cKO by refill mechanisms, which kept up the necessary pool size to produce astrocytes, but not upper layer neurons.

### DOT1L affects astrogenesis from ventral progenitors

A fraction of cortical astrocytes is described to derive from *Nkx2.1*-expressing progenitors, residing in the ventral telencephalon [11]. Given that DOT1L impacts adaptation of specific cell fates in stem cells in diverse tissues [28,51,52], we also analysed a *Nkx2.1*-cre DOT1L LOF by multiome snRNA-seq, specifically in the dorsal telencephalon at E16.5 (**Fig. 8a**) for assessing DOT1L’s impact on astrocytes in this setting. Notably, we expected *Nkx2.1-cre* astrocytes to be a minor fraction compared to NSC/AP-derived astrocytes in the dorsal telencephalon. We identified Early Immature Astrocytes, and RGC with astrocytic gene expression signature in the *Nkx2.1-cre* DOT1L LOF data set from their respective marker gene expression profile (**Fig. S5**). Surprisingly, cell proportions analyses revealed a slightly higher fraction of Early Immature Astrocytes at E16.5 upon DOT1L LOF in *Nkx2.1*-expressing cells (**Fig. 8b**). Both astrocyte populations had a set of significant DEGs upon DOT1L LOF (**Fig. 8c**), and the smaller proportion of *Nkx2.1*-derived astrocytes in the entire population of cortical astrocytes might explain the relatively small number of significant DEGs in this *Nkx2.1-cre* derived DOT1L LOF. Comparison of significant DEGs in cortical astrocyte populations in the P0 *Emx1-cre* (WT and cKO combined) and E16.5 *Nkx2.1-cre* data sets revealed, as expected, few shared DEGs. Comparing DEGs from E16.5 *Nkx2.1*-derived astrocyte populations with significant DEGs from Astrocytes I and II from P0 indicated a higher number of overlapping genes with Astrocytes I compared to II (**Fig. 8d**). Comparing within both E16.5 data sets revealed the highest overlap between Early Immature Astrocytes (Nkx2.1 data) and Early Astrocytes (Emx1 data). Functional GO term enrichment analysis retrieved that within the Nkx2.1 data, both astrocyte populations reduced expression of translational genes (**Fig. 8e**), similar to Astrocyte I features, and in accordance with the DEGs overlap. *Emx1-cre* DOT1L LOF instead increased translation of genes in Early Astrocytes, featuring an Astrocytes II phenotype. This finding was in accordance with an increased contribution of Astrocytes II to Early Astrocytes between E14.5 and E16.5 in *Emx1-cre* Dot1l cKO compared to WT (**Fig. 5c, d**). As specific TFs impacted astrogenesis at E16.5 (**Fig. 6a, b**), we also determined DEGs among the Early Immature Astrocytes in the Nkx2.1 data, showing *Lhx2* with decreased expression, similar to Emx1 data (**Fig. 6a, 8f**). Cell oracle perturbation of *Lhx2* in Nkx2.1 data led to an opposite effect compared to Emx1 data, because the knock down of *Lhx2* was predicted to promote the Early Immature Astrocyte state over the RGC/Astrocytes, whereas the overexpression in the cKO favoured the RGC/Astrocytes. This was in accordance with the cell proportion changes (**Fig. 8b**).

**Figure 8.**
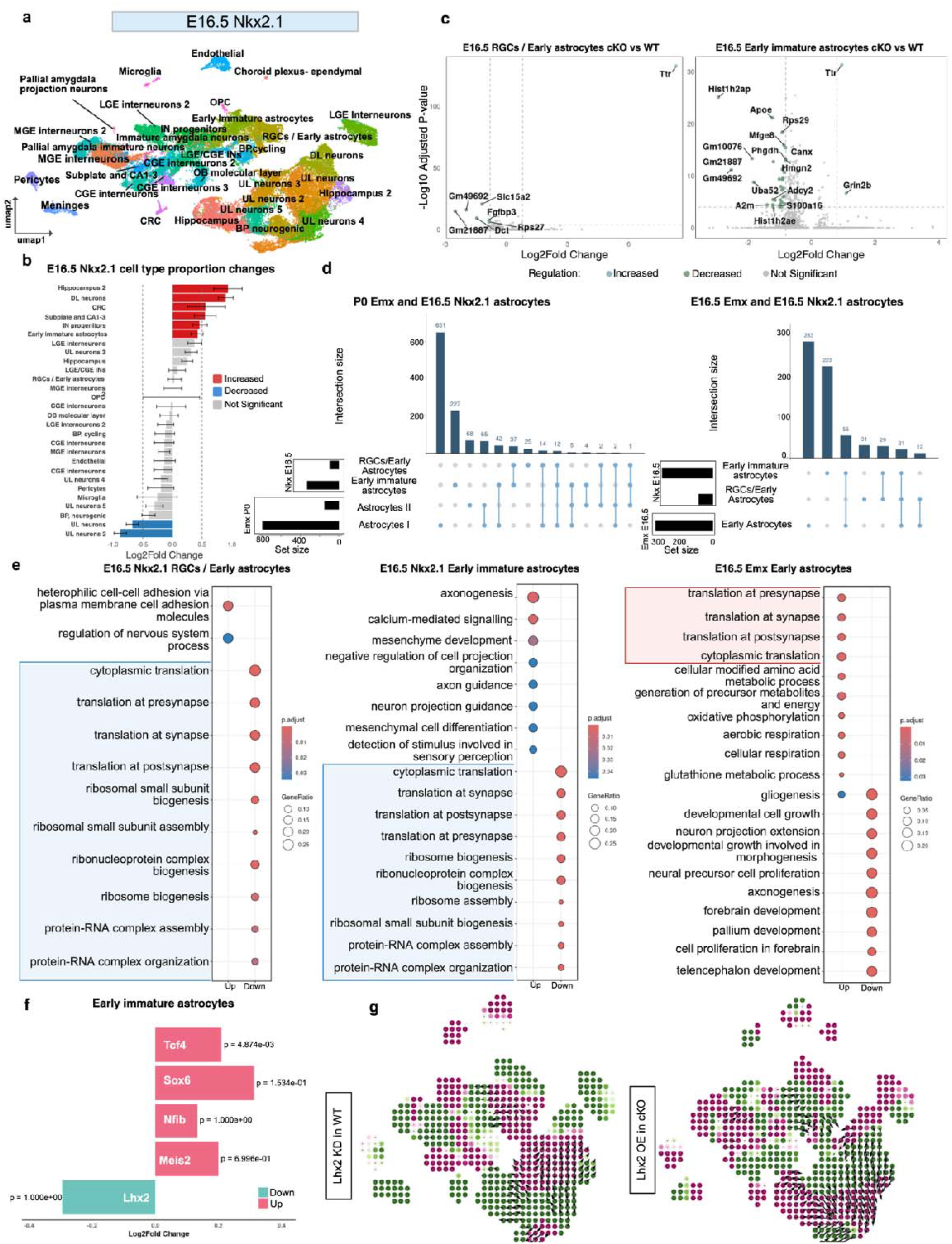
**a** UMAP representation of E16.5 (left) Nkx2.1 datasets. Colour represents different clusters. **b** Bar plot illustrating cell proportion changes upon Dot1l cKO. Significantly increased or decreased cell types were defined with thresholds FDR < 0.05 and Log2Fold Change > 0.40. Error bars represent 95% confidence intervals from 1000 permutations. Red indicates significant increase, blue significant decrease, and grey no significant change. **c** Volcano plot depicts the increased (blue), decreased (green) and not significant genes (grey) for *Nkx2.1-cre* E16.5 RGCs/ Early astrocytes cKO vs WT (left), and *Nkx2.1-cre* E16.5 Early immature astrocytes cKO vs WT (right). Horizontal dashed line shows where p-value is 0.05, and the vertical dashed lines show where the fold change is 1 and −1. **d** Upset plot showing the overlap of DEGs between *Nkx2.1-cre* E16.5 (RGCs/ Early astrocytes and Early immature astrocytes) and *Emx1-cre* P0 Astrocytes I and II (left) and between *Nkx2.1-cre* E16.5 (RGCs/ Early astrocytes and Early immature astrocytes) and *Emx1-cre* E16.5 (Early Astrocytes) (right). **e** GO biological process enrichment analysis of significant DEGs upon *Nkx2.1-cre* Dot1l cKO for E16.5 RGCs/ Early astrocytes (left), E16.5 Early immature astrocytes (middle), and *Emx1-cre* E16.5 Early Astrocytes (right). Significantly enriched translation and biogenesis processes associated with increased (right red box) and decreased gene expressions (right blue boxes). Enrichment was performed with Benjamini–Hochberg correction (*p* < 0.05, *q* < 0.2). **f** TFs changing upon *Nkx2.1-cre* Dot1l cKO in Early immature astrocytes. Change is shown using Log2Fold Change and P-value significance is shown on the bars. Green and pink colours denote decrease and increase upon the Dot1l cKO respectively. **g** Simulation vectors show the predicted transition in cell states after *Lhx2* KD in WT (left) and OE (right) in E16.5 *Nkx2.1-cre* Dot1l cKO datasets representing the altered likelihood of a cell state change post perturbation. Pink and green denote the effects of perturbation on lineage trajectories. Pink indicates a deviation from the expected developmental trajectory, whereas green indicates that the differentiation trajectory remains aligned with the expected developmental lineage. The arrows represent the predicted shift in cell identity in response to the TF perturbation. KD: knockdown, OE: overexpression

Together this data shows that DOT1L regulates similar expression programs within two different astrocyte lineages, i.e. *Emx-cre* and *Nkx2.1-cre* derived, by controlling a set of instructive TF. Notably, this set of TF is not only active during astrogliogenesis, but also during neurogenesis.

## Discussion

Our study provides refined views on cortical astrocyte development. First, we highlight prenatal expression programs in astrocytes that are highly diverse concomitant with transcriptional plasticity. We also show diverse origins of astrocytes. In this context, we challenge the prevailing view that APs are the only source of cortical astrocytes by showing that a substantial fraction of astrocytes have an origin from NSCs, especially in early cortical development. Further, we refine the view that astrogenesis strictly follows neurogenesis, as APs feed into the astrocyte lineage at the earliest time point, we analysed, i.e. E12.5 during the peak of neurogenic proliferation. Our data are further substantiating that cortical astrocytes also derive from the *Nkx2.1* lineage, with progenitors residing in the ventral telencephalon. In addition, we exposed the epigenetic layer controlling astrocyte development by placing this study in the context of DOT1L LOF, mediating H3K79 methylation and adjusting transcriptional programs. Essentially, we expose that the plasticity of astrocytes to convert from mature states to progenitors, including NSCs and APs, occurs in WT conditions, but expression of SOX2-positive stem cells features is increased in astrocytes upon DOT1L inhibition or LOF. Lastly, our study highlights a set of TFs that are at play during astrogenesis and state conversions, including TF also impacting neurogenesis. This indicates that the assumed switch from neuro- to astrogliogenesis will be accompanied by gross adaptation of the chromatin landscape that allows activity of similar TF to drive transcriptional programs that are so diverse that either neurons or astrocytes are the phenotypic end product. In this regard, DOT1L drives astrogliogenesis by regulating a TF network, including *Nfib, Meis2*, and *Lhx2*, and regulates astrocyte state conversion through *Nfib, Meis2, Sox6*, and *Tcf4*.

### Plasticity of transcriptional programs in embryonic astrocytes

A recent report described a dual origin of astrocytes in regard to their expression of *Emx1, Olig2* and *S100a11* [9]. Following up on these results and using marker gene suggestions, we assessed our data for similar signatures. However, neither our own data, nor enriched data sets from publicly available brain cell atlas studies [40,41] allowed us to use the same top enriched transcriptional gene sets for characterising embryonic astrocyte populations. Assumingly, embryonic astrocytes are a heterogeneous class of cells that convert between progenitor and mature stages, even towards stem or progenitor cells like NSCs or APs. This restricts their classification based on a set of established marker genes including *Aldh1l1, Aldoc, Ndrg2, Apoe*, and *Slc1a3*. These markers, however, are also active to certain extent in RGCs. For correct assignment of astrocytes especially in single-cell data sets, an enlarged marker set is desirable, of which our study exposed a potential set that together with the canonical known markers might strengthen annotations during embryonic development.

The transcriptional plasticity of early astrocytes, allowing to re-activate stem cell programs or adaptation of mature phenotypes, might serve a necessary flexibility to adapt to specific needs of the developing cerebral cortex. Given individual differences for example in the timing of neurogenesis, it might be important for the astrocytic niche to adapt to a different appearance or maturation of a neuronal network, in mice equally as humans [53]. Similarly, time and spatial variation in the emerging blood vessels [54] might also require flexible astrocyte states while building the blood-brain-barrier.

Cell lineage plasticity bears important insights, as it might be the basis for the observation that conditional mutants often retain a set of cells that should have been targeted. For example, if one assumes *Emx1* to be expressed in all APs, all progeny should bear a cKO if *Emx1-cre* is used. However, nearly all conditional mutants, including ours, have a fraction of resilient cells that behave normally. Our data suggest that partial replenishment of stem cell pools from other progenitor populations is one mechanism driving resilience towards “lineage-specific” cKOs.

### NSCs contribute to diverse origins of embryonic astrocytes

Highly represented in literature on cortical development is the notion that APs first generate neurons, switching to astrogliogenesis later in development. This can be interpreted in a way that APs are the only source of astrocytes. Our data show that this is not the case, as NSCs are an additional source of astrocytes in the cortex, and ventral progenitors contribute as well [11]. Our data also suggest that APs produce astrocytes during the neurogenic phase of development. It is thus possible that mechanisms exist that mediate an early switch towards astrogliogenesis. An equal explanation could be that asymmetric divisions include production of daughter cells, one with the competence to generate astrocytes while the others stay in the neurogenic lineage. Exploring such potential heterogeneity of NSCs and APs with corresponding high resolution is tempting for future experimentation. Interestingly in regard to the plasticity of stem cells during cortical development, is that astrocytes can de-differentiate towards NSCs and APs. Reactivation of stem cell programs in astrocytes is a critical point in reprogramming towards neurons. In this regard, it has been reported that overexpression *of Oct4, Sox2, Nanog* [13] or *Dlx2* [14] led to *Ascl1*-expressing neural progenitor cells. Strikingly, our data suggest that this de-differentiation occurs naturally in development, with the most prominent conversions from Astrocytes I towards APs III, Astrocytes II to uncommitted NSCs, and Astrocytes III to APs II. Not only is the insight from our study useful to refine reprogramming attempts, i.e. by inhibition of DOT1L activity, but it might also reflect adjustments for individual differences in development.

Our findings can also be placed into the context that ablation of progenitors can be compensated by clonal expansion of resilient progenitors. Recent observations suggest that following an early reduction of the cortical progenitor pool in development, the neuronal output recovers through compensatory refill mechanisms, by which clonal amplification of specific progenitors restores the missing fraction [55]. We hypothesise, based on the finding of an *Emx1*-negative origin of astrocytes, that an early DOT1L LOF could bias towards reducing or losing *Emx1*-positive progenitors. An increased fraction of *Emx1*-negative APs from equally negative NSCs would be the source to compensate for the DOT1L-LOF mediated loss of astrocytes, but maybe not for the neurogenic cell lineages. Going beyond cellular phenotypes, it is conceivable that the genome of contributing cells should accordingly have epigenetic plasticity for activating necessary transcriptional adaptations. DOT1L preserves cell states in the CNS [26-28,56], and our data presented here are expanding this view extending from neurogenesis to astrogliogenesis. As DOT1L inhibition also increases reprogramming efforts towards stem cells from somatic cells [57,58], our finding of increased expression of the stem cell marker SOX2 upon DOT1L inhibition emphasises that cell state and fate switches can be facilitated upon DOT1L LOF. As technical advancement on the single cell level shall allow determining altered epigenomes beyond scATAC, including histone modifications and TF distribution, these experiments will probably expose many more insights into coupling cell and epigenomic plasticity in the future.

### A set of TF regulates astrogliogenesis and astrocyte state conversion

Our data also refine common views, upon which astrocytes are first detected in the mouse cerebral cortex at E16.5 [59]. While taking the single entities of our RNA-seq data at E12.5, E14.5 and E16.5 into account, our data are seemingly in support of this finding. However, integration over multiple time points and including extending data sets, identified astrocytes also at earlier time points. We used the *Emx1-cre* and *Nkx2.1-cre* dependent deletion of DOT1L exon 2 to confirm a heterogeneous origin of astrocytes, corroborating reports on multiple origins of cortical astrocytes, for example, from *Emx1*- and *Olig2*-expressing progenitor lineages [9], as well as from *Nkx2.1*-expressing ventral progenitors [11]. Diverse progenitors defining the origin of astrocytes can be one source of heterogeneity that can perpetuate into distinct molecular features of these cells in different brain regions [60]. The activity of distinct TFs, including *Emx1* and *Olig2* for cortical astrocytes [9], or *Nkx2.1* and *Zic4* for septal astrocytes [61] might be implicated to establish specific molecular features. Our data suggest that an extended network of TFs regulates astrogliogenesis and astrocyte state conversion. Our interpretations are based mainly on bioinformatics prediction, and the precise roles for TFs need to be confirmed experimentally in future attempts in suitable model systems. For example, we predict a role for TCF4 in keeping mature astrocyte states. However, till now TCF4 is reported to prevent astrocyte fate in the *Nkx2.1* lineage. i.e. astrocytes derived from ventral progenitors. Extending this view, *Tcf4* reduction fosters generation of Early Astrocytes from both APs and BPs [20]. However, the impact of *Tcf4* on astrogliogenesis might be regionally restricted in the dorsal telencephalon, i.e. to midline glia [45]. We aimed to resolve the impact of *Tcf4* expression on astrogenesis in more detail; however, the public data set [42] providing scRNA-seq data only contains few astrocytes in the LOF condition, hampering the deduction of quantitative changes in cell proportions. We used *in silico* perturbations to survey the impact of specific TF on progenitor (AP/BP) fates towards astrocytes, and whether differences would produce different states of astrocytes that confer phagocytosis or homeostasis. After perturbation of the activity of specific TFs *in silico*, we revealed at P0 that *Tcf4* and *Nfib* regulated expression programs favouring AP to Astrocyte II differentiation. *Nfia*, is known to favour astrogenesis [62], and our predictions are in support of this notion, as it impinged on AP to Astrocyte I generation. *Meis2*, probably in cooperation with *Nfib*, drives astrocyte differentiation from APs at E16.5. *Sox6* and *Lhx2* promote astrogliogenesis as well [62], a finding that is also corroborated by our data. Thus, despite the fact that our predictions are limited by being based on *in silico* analyses using one bioinformatics tool, published experimental data support our exemplarily interpretations of instructive TF networks during astrocyte generation and differentiation.

### Epigenetic processes as regulative layer of astrogenesis

Notably, the TF mentioned above and at play during astrogliogenesis are also known to impact neurogenesis, thus their cell type specific functions depend on further factors, including the local environment and signalling pathway activities, e.g. SHH or FGF for astrocytes within the septal regions [61]. Our data expose the epigenetic landscape as a further layer of molecular specification of astrocyte function. While this study did not delve into the detailed description of single-cell resolved chromatin features underlying the switch from neuro- to astrogliogenesis, the extension from DNA methylation as regulatory switch to the activity of a specific histone methyltransferase, DOT1L, opens up a new perspective linking the epigenome to astrocyte’s development. Whereas our *in vivo* data expose a role for DOT1L in control of astrogliogenesis and astrocyte functional state conversion, it was recently reported that mouse embryonic stem cells (mESC) bereft of DOT1L exon 5 were incapable of activating glial expression programs whereas neurogenesis was unaffected [63]. This finding is seemingly at odds with the here presented observations and other reports on DOT1L functions *in vivo*, in which both neuro- and astrogliogenesis appear impaired, but not confined to either one of the two cell lineages. This contradiction is probably resolved considering that the DOT1L LOF at mESC stage is much more severe compared to its deletion after initial ectodermal priming and execution of primary NSC/NPC development, where DOT1L conserves cellular states and adapts transcriptional programs. Among the DOT1L-regulated TF that affect astrogliogenesis we identified *Lhx2*, which is reported to limit astrogliogenesis in the developing hippocampus [46] and cortex [16]. However, its function might be context-dependent, as it is necessary for the development of retinal astrocytes [64], and has been identified as an astrocyte-promoting factor in mESCs (supplement of [62]). Further TFs under control of DOT1L which affect astrocyte development positively, we identified *Nfia* [62], *Sox6* [62], and *Zbtb20* [65]. *Creb5* expression correlates with regional differences of cortical and thalamic astrocytes [66], and it impacts astrocyte metabolism [67], however, functional data about astrocyte development are sparse. Our cell oracle prediction supports a role of *Creb5* in shifting Early Astrocytes towards APs upon overexpression in cKO cells at E16.5, suggesting a role for this TF in astrocyte development as well.

Taken together, this study refines important concepts of astrocyte development during cortical development by challenging common views on timing and stem cell contribution in astrogliogenesis. Our work also highlights the necessity to explore the epigenetic regulation at the interface of neuro- and astrogliogenesis, and benchmarking epigenetic contribution by exploring DOT1L function. Nonetheless, further experimentation is needed to resolve at the single-cell level the epigenetic traits involved and responsible for lineage specification decisions and associated plasticity.

## Data availability

Raw scRNA sequencing data from E12.5, E14.5, E16.5 and P0 cortex for both Emx1-cre Dot1l cKO and WT samples and from E16.5 Nkx2.1-Cre KO and WT samples, have been deposited in the Gene Expression Omnibus (GEO) and will be made available upon publication. LaManno scRNA-seq data is available at the Sequence Read Archive (SRA) under accession number: <u>PRJNA637987</u>. The DiBella mouse dataset used for this study is available at <u>GSE153164</u>. Raw data for the E18.5 scRNA-seq of Tcf4-cKO mice is accessible via: <u>GSE147247</u>.

## Code availability

All Python and R Markdown scripts and conda environments used to generate the main and supplementary figures are available on GitHub: <u>DOT1L-Effect-In-Astrogenesis</u>. Objects to reproduce the figures are stored in Zenodo and will be made accessible upon publication.

## Acknowledgment

We sincerely thank Sebastian Preissl, Institute of Experimental and Clinical Pharmacology and Toxicology, University of Freiburg, Germany, for providing access to his laboratory for library preparation. We are grateful to Arquimedes Cheffer, Institute of Anatomy and Cell Biology, University of Freiburg, Germany, for valuable discussions and assistance with wet lab experiments. We also thank Ulrike Bönisch, Max-Planck-Institute of Immunobiology and Epigenetics, Freiburg, Germany, from the sequencing core facility. We also appreciate the bioinformatics core facility at the MPI Immunobiology and Epigenetics for their support and feedback with the data analysis. This study was supported by the German Research Foundation 322977937/GRK2344 (TM, TV).

## Author’s contribution

The authors have no relevant financial or non-financial interests to disclose.

TM and TV conceived and designed the study. CLF was responsible for library preparation, and data collection. MM performed the data analysis. CV conducted the other wet-lab experiments. TH and JR contributed to conceptualization, quality control of the analysis. The first draft of the manuscript was written by TV, and figures prepared by MM and TV. All authors read, edited and approved the final manuscript.

