## Supplementary figures and table for "Epigenetic perturbation unravels diverse origins, state conversions and de-differentiation of cortical astrocytes"

### *Supplementary 1*

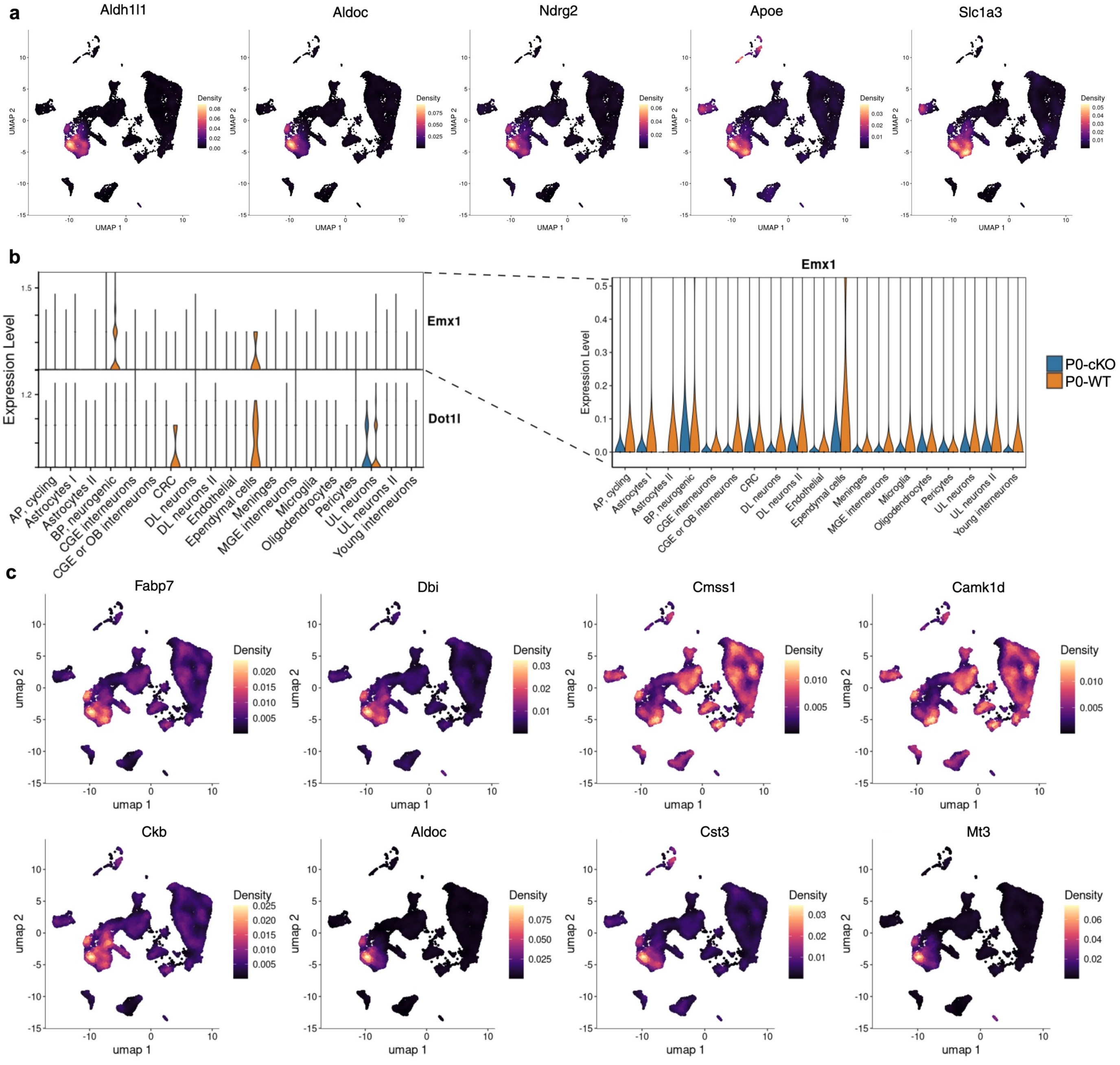


**1. *Astrocyte marker genes, Emx1 and Dot1l expression at P0 in control and DOT1L LOF cortex***

**a** Density plot illustrating expression of canonical astrocyte marker genes in P0 dataset. **b** Violin plot showing the *Emx1* and *Dot1l* expression across all P0 cell types (left). Zoomed in *Emx1* expression across cell types. **c** Density plot showing the enriched differentially expressed genes in Astrocyte II population.

### Supplementary 2

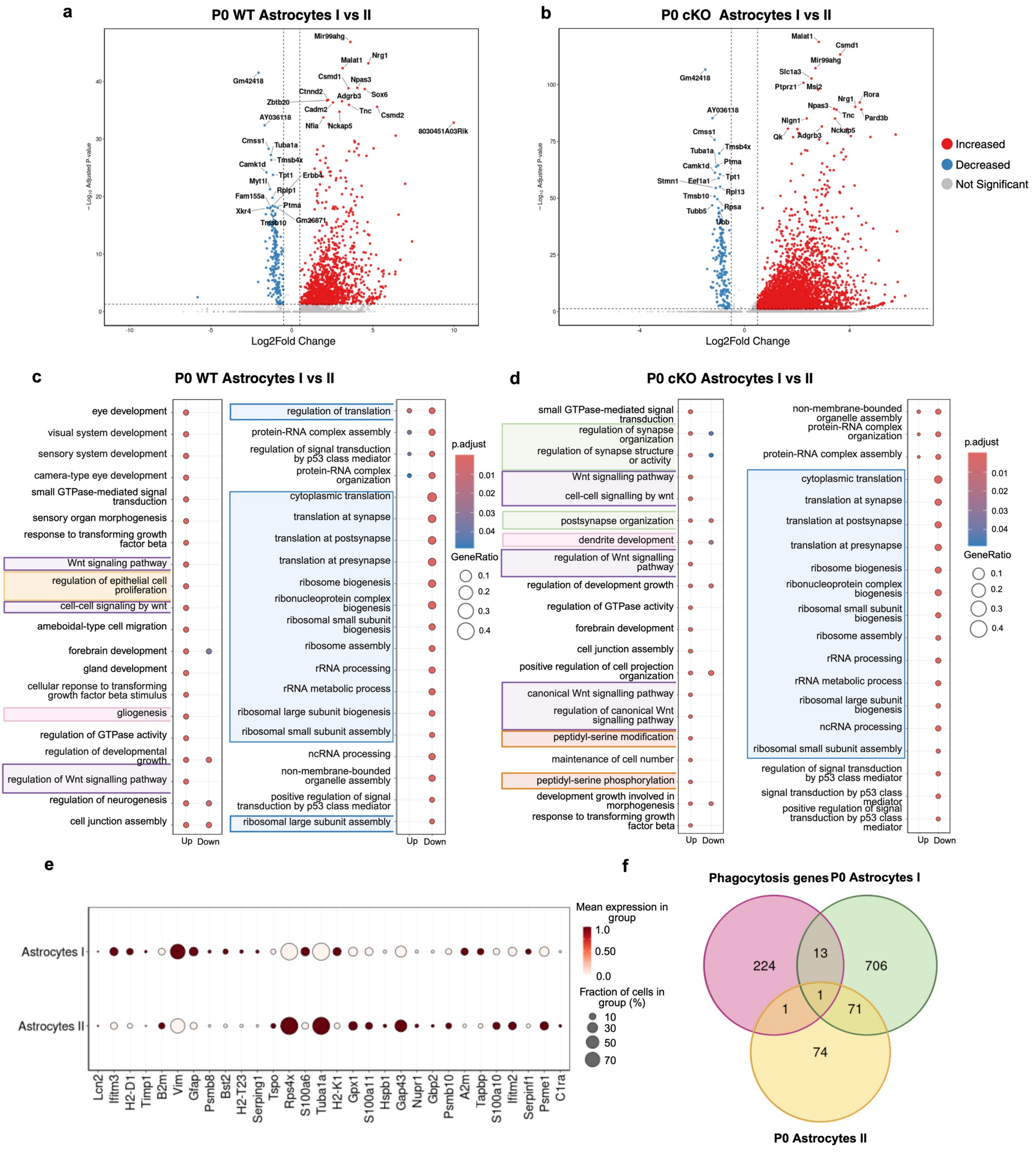

**2.** ***DOT1L limits synaptic functions and serine modification in Astrocytes I compared to Astrocytes II***

**a** Volcano plot of genes that increased (red) or decreased (blue) significantly in P0 WT Astrocytes I vs Astrocytes II comparison, and **b** in P0 cKO Astrocytes I vs Astrocytes II. Horizontal dashed line indicates the significance threshold of adjusted p-value = 0.05, and the vertical dashed lines show the Log2Fold Change threshold at 0.5 and -0.5. **c** GO term enrichment analysis of significant DEGs between P0 WT Astrocytes I vs Astrocytes II. **d** GO term enrichment analysis of significant DEGs between P0 cKO Astrocytes I vs Astrocytes II. Significantly enriched biological processes associated with different groups are indicated by color: green – synaptic processes, purple – Wnt associated pathways, orange – peptidyl-serine related processes, pink – axon-related and neurodevelopmental, yellow – cell proliferation and blue translation and biogenesis related processes. Enrichment in e and f was performed using Benjamini–Hochberg correction ($p < 0.05$, $q < 0.2$). **e** Dot plot showing the expression of reactive astrocyte marker genes across Astrocyte I and II populations. **f** Venn diagram illustrating the overlap between significant DEGs from P0 Astrocyte I and II populations and phagocytosis-associated genes obtained from GO term GO:0006909.

### Supplementary 3

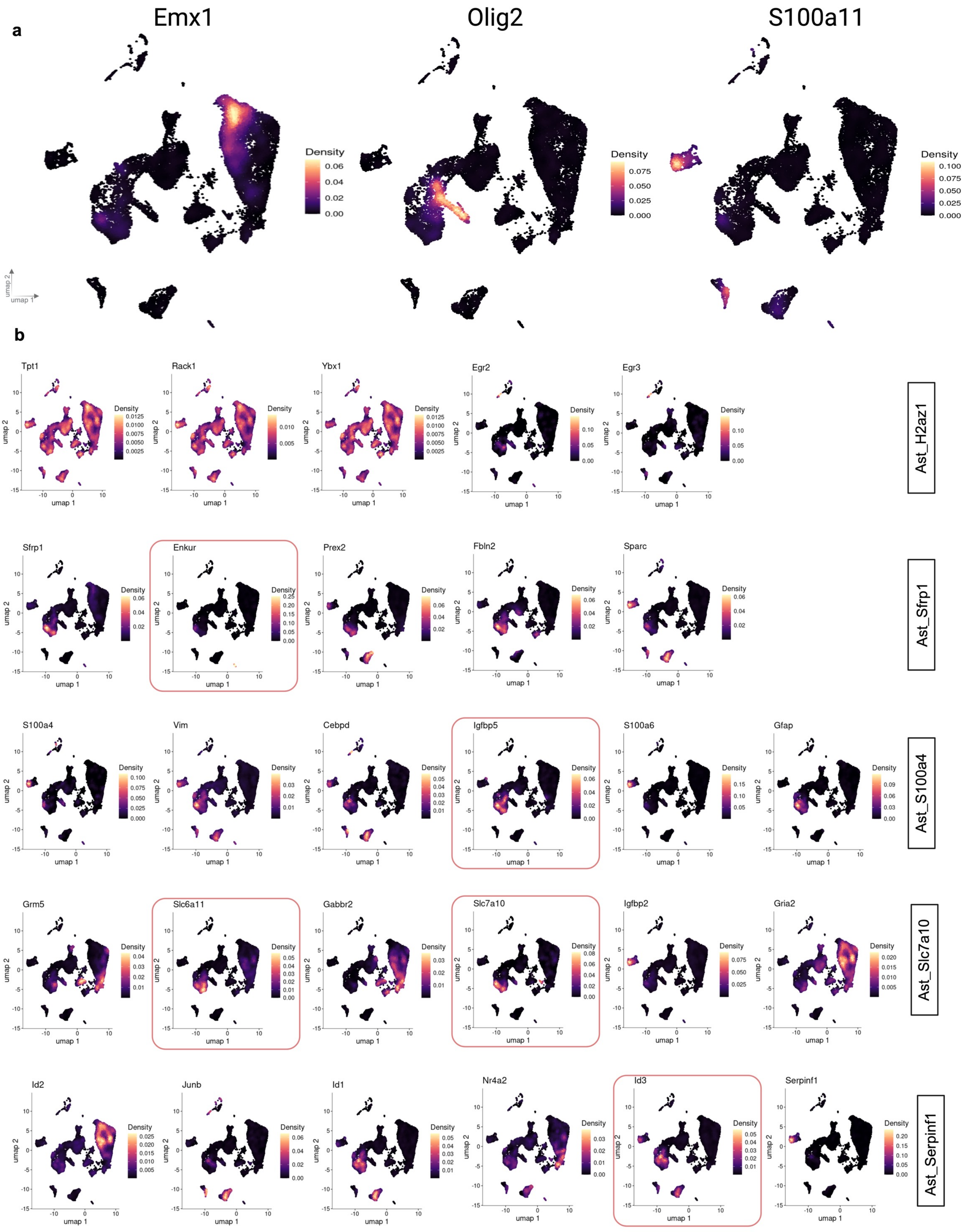

**3. *Expression of embryonic and postnatal astrocyte marker genes in P0 astrocytes***

**a** Density plots illustrating the expression levels of *Emx1*, *Olig2* and *S100a11* across cell types in P0 dataset (genotypes combined). **b** Density plots illustrating the expression of genes of different astrocyte subtypes, described in [9] and named accordingly as indicated on the right, in P0 combined data. Red squares highlight common genes with expression in Early Astrocytes at E16.5, and Astrocytes I and II at P0.

### *Supplementary 4*

a

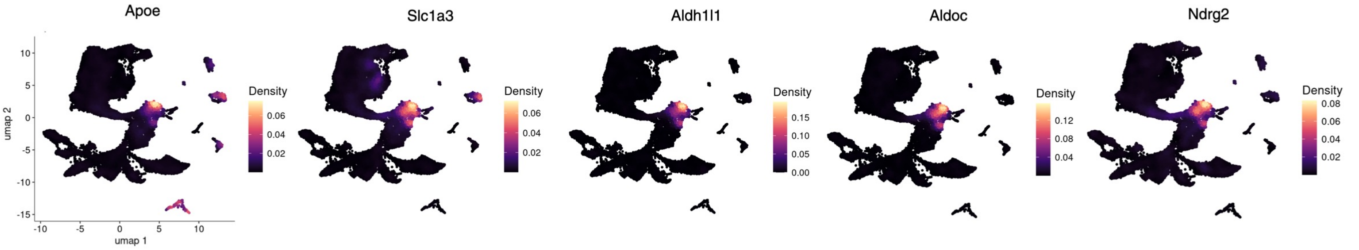


b

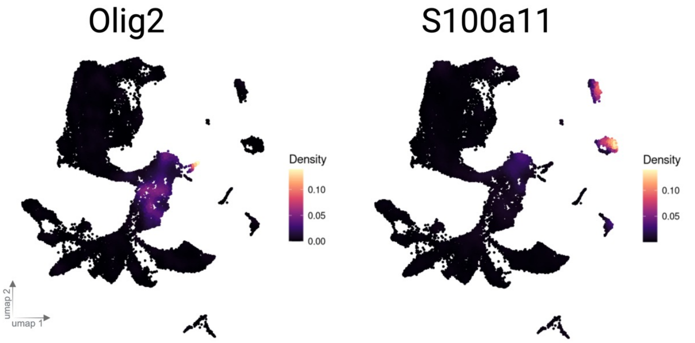


c

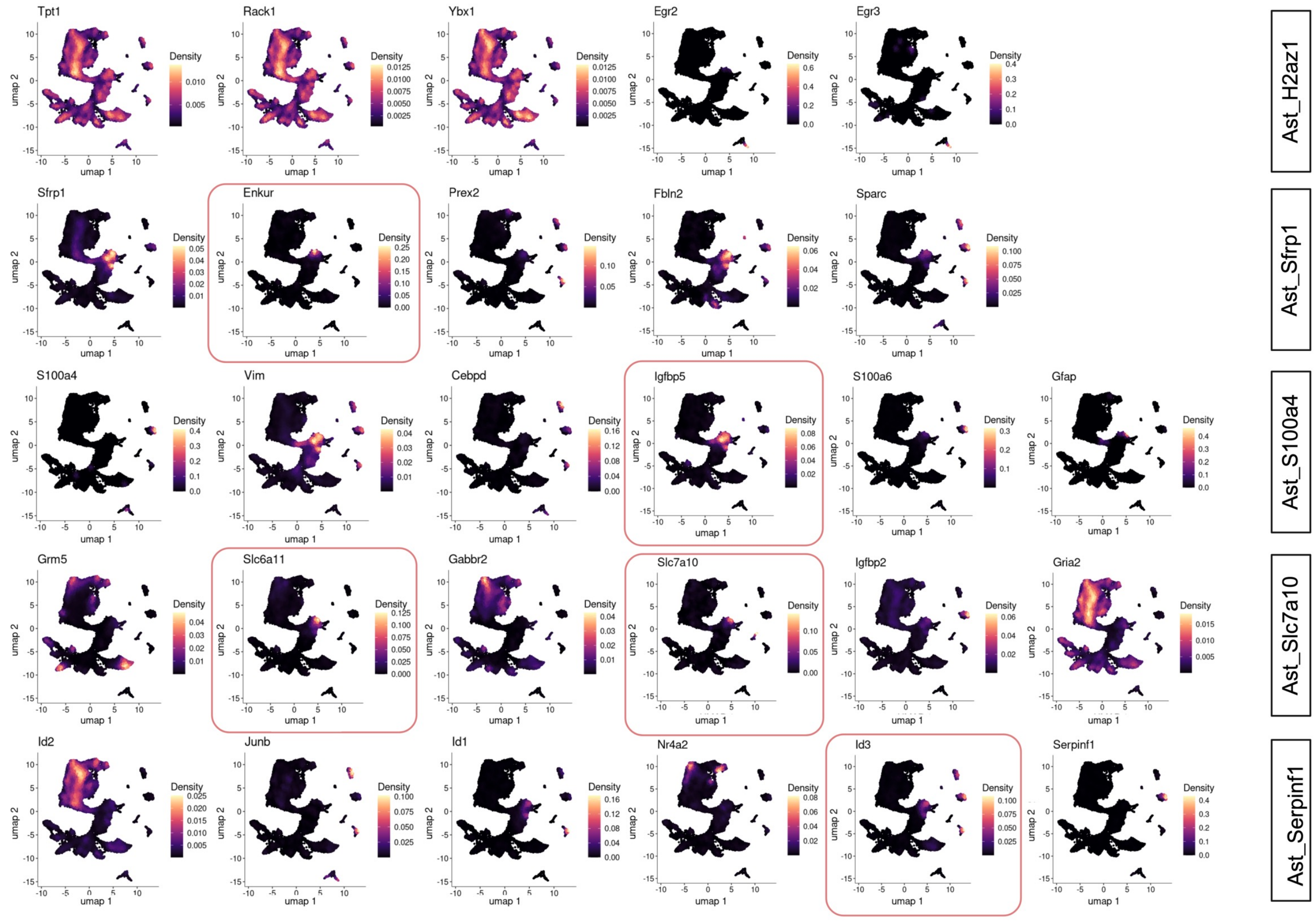

**4.** ***Expression of astrocyte marker genes in E16.5 astrocytes***

**a** Density plot illustrating expression of canonical astrocyte markers in the E16.5 dataset. **b** Density plots showing the expression levels of *Olig2* and *S100a11* across cell types in the E16.5 dataset. **c** Density plots illustrating the expression of genes of different astrocyte subtypes, as described in [9], at E16.5. Red square highlights the genes that are expressed within the Early Astrocyte cluster.

### Supplementary 5

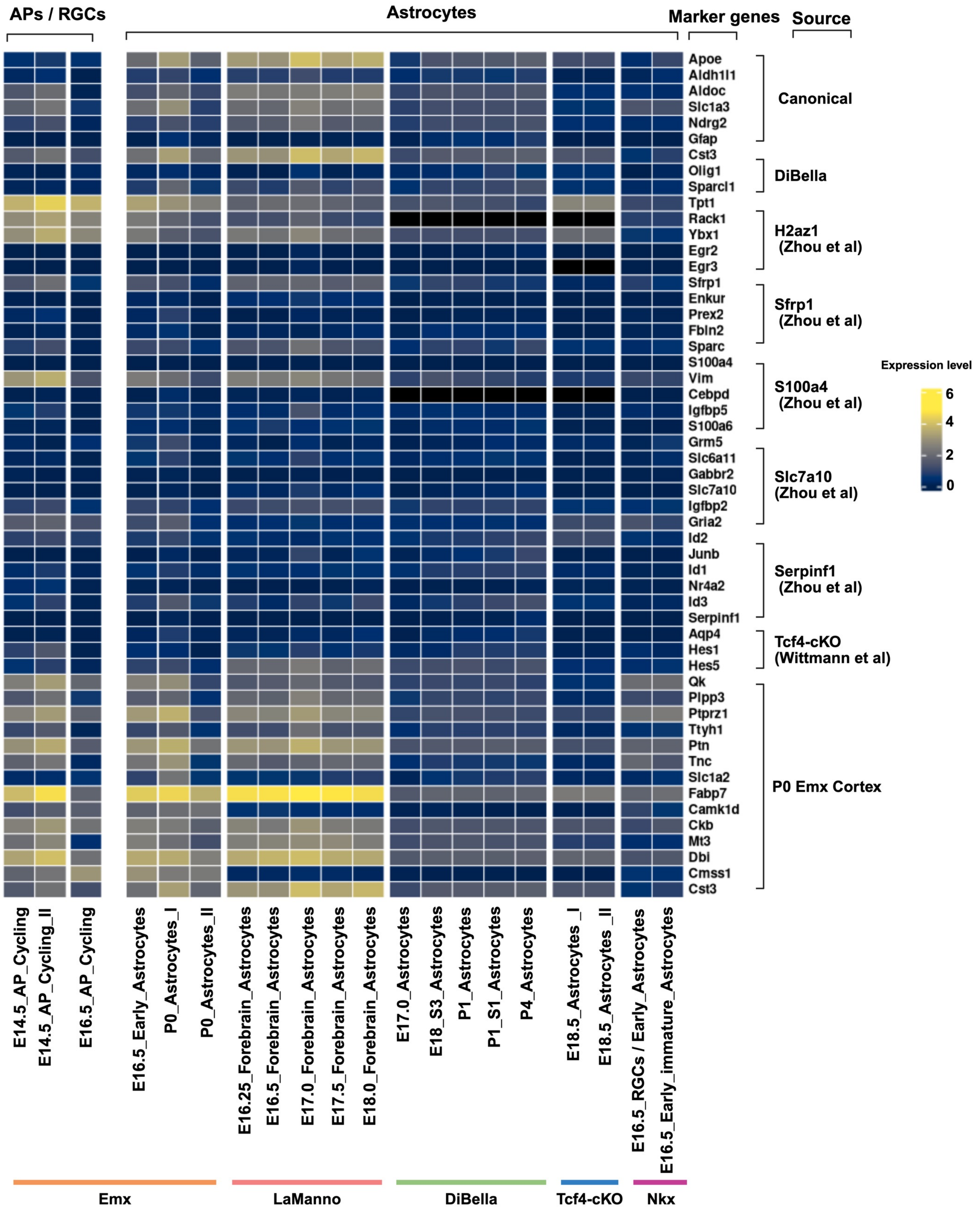

**5. *Marker gene classification in embryonic, pre- and postnatal astrocytes from various data sets reveals a set of genes characterising embryonic astrocytes***

**a** Heatmap of astrocyte marker gene expression across multiple datasets (Emx (this study), La Manno [40], Di Bella [41], Tcf4-cKO [42], Nkx (this study)) covering APs/RGCs and astrocytes. On the Y axis marker gene names are given for which the expression is shown, together with source name in which these marker genes appear as top enriched in the respective astrocyte population. On the X axis cell types expressing the marker genes are shown with their timepoint indicated. Sources of the cell types/datasets are highlighted with colors (Emx – orange, LaManno – red, DiBella- green, Tcf4-cKO – blue and Nkx – pink). APs: Apical Progenitors, RGCs: Radial glial cells.

### *Supplementary 6*

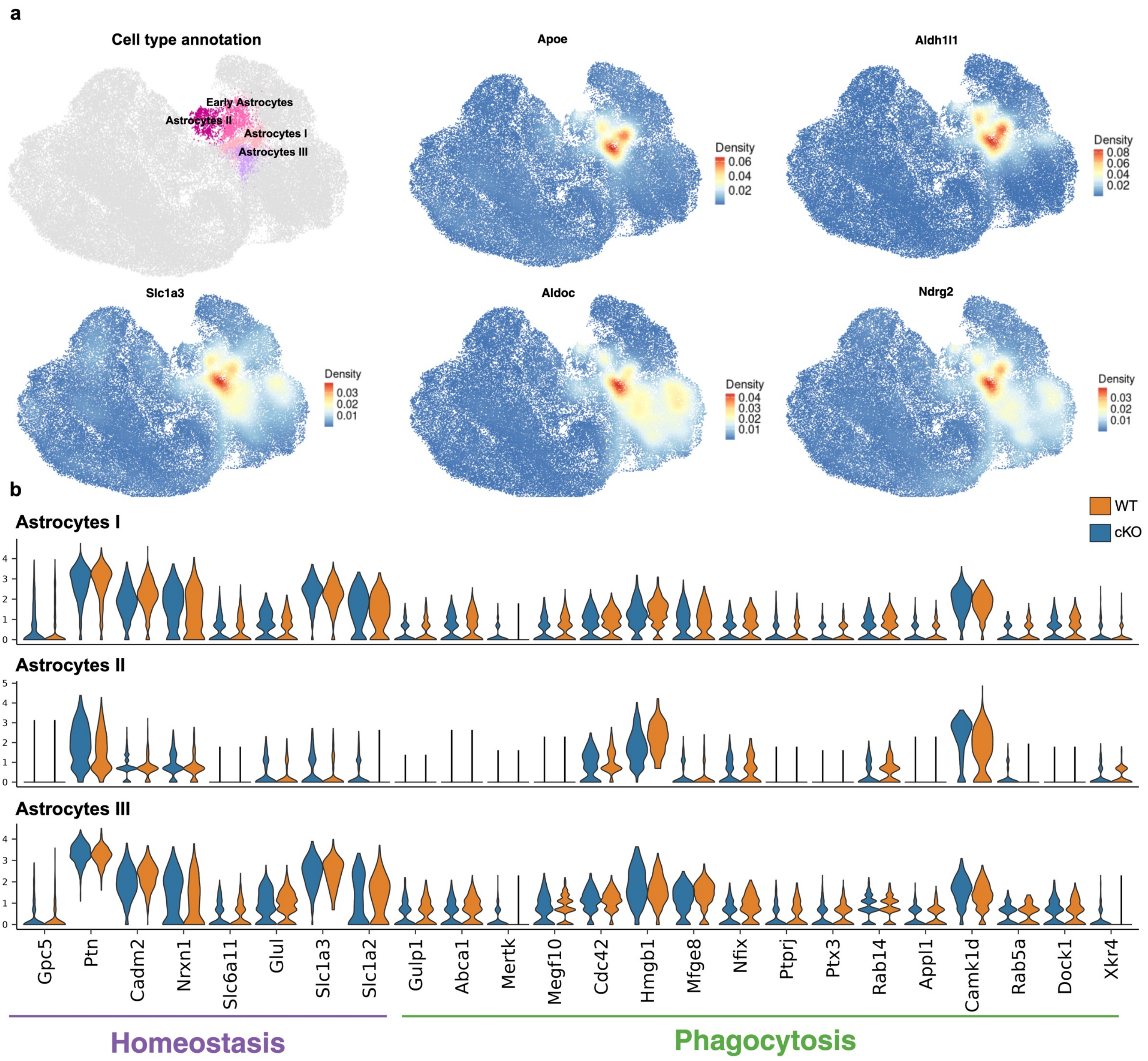

### Supplementary 6

c

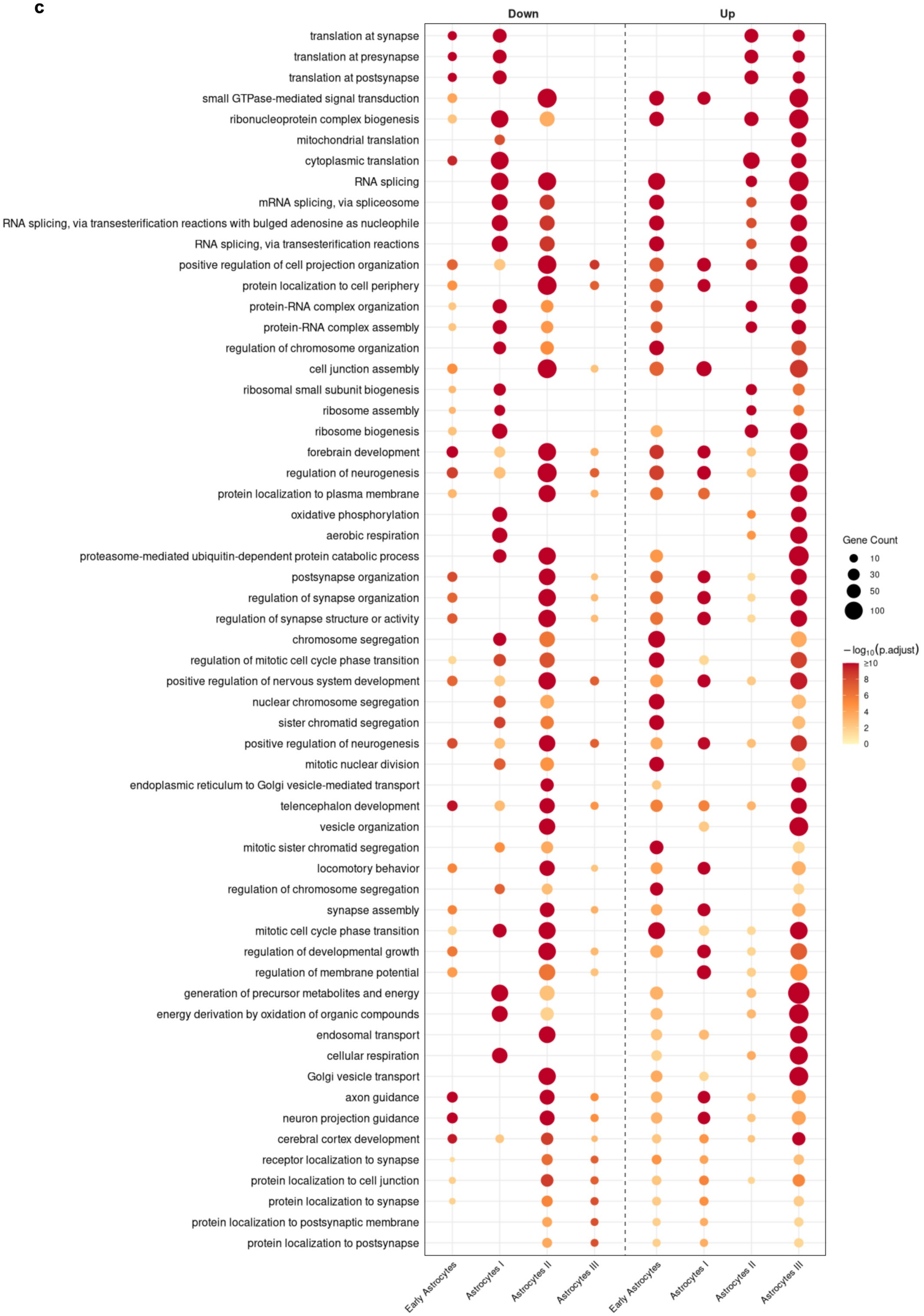

**6.** ***Gene expression in different astrocyte clusters upon data integration over embryonic and postnatal time points***

**a** Density plot illustrating the expression of canonical astrocyte marker genes in the integrated dataset (E12.5–P0). **b** Violin plots showing phagocytosis and homeostasis marker gene expressions across different astrocyte clusters (Astrocytes I-III) and split according to genotype. **c** GO term enrichment analysis of significant DEGs across different astrocytes (Early Astrocytes, Astrocytes I-III) populations. Biological processes going down (left) and up (right) separated by a dotted line. Enrichment analysis was performed with Benjamini–Hochberg correction ($p < 0.05$, $q < 0.2$).

### *Supplementary 7*

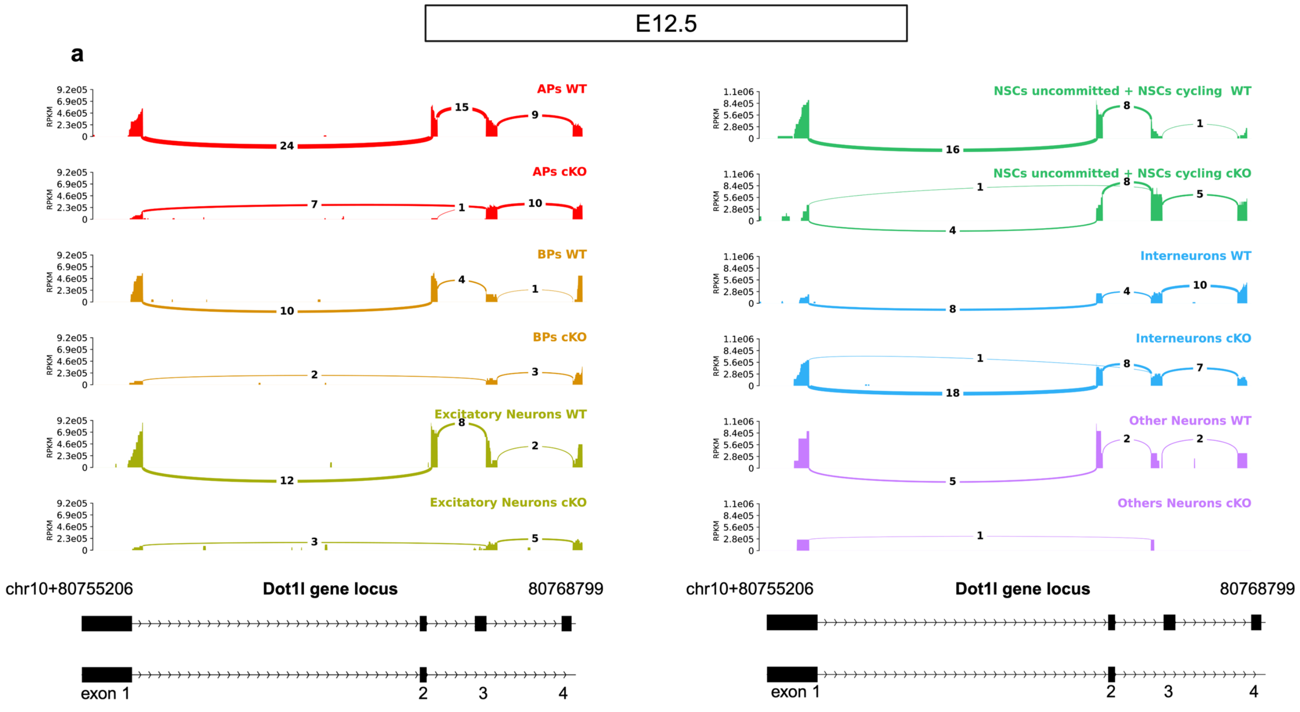


**7. *Emx1-cre activity in different cell populations at E12.5 supports lineage trajectories based on presence of Dot1l exon 2***

**a** Sashimi plots displaying the RNA-seq read coverage (RPKM) and splice junction usage at the *Dot1l* gene locus (chr10: 80,755,206–80,768,799) in E12.5 excitatory neuron lineage cell types (left), and other E12.5 neural cell types (right). Filled areas represent per-base read depth scaled in RPKM. Arcs connecting exonic regions represent split reads spanning splice junctions, with the number above each arc indicating the number of junction-supporting reads. The gene model at the bottom depicts annotated *Dot1l*, with black boxes representing exons and horizontal arrows indicating intron direction on the forward (+) strand. Exons 1–4 are labelled.

### *Supplementary 8*

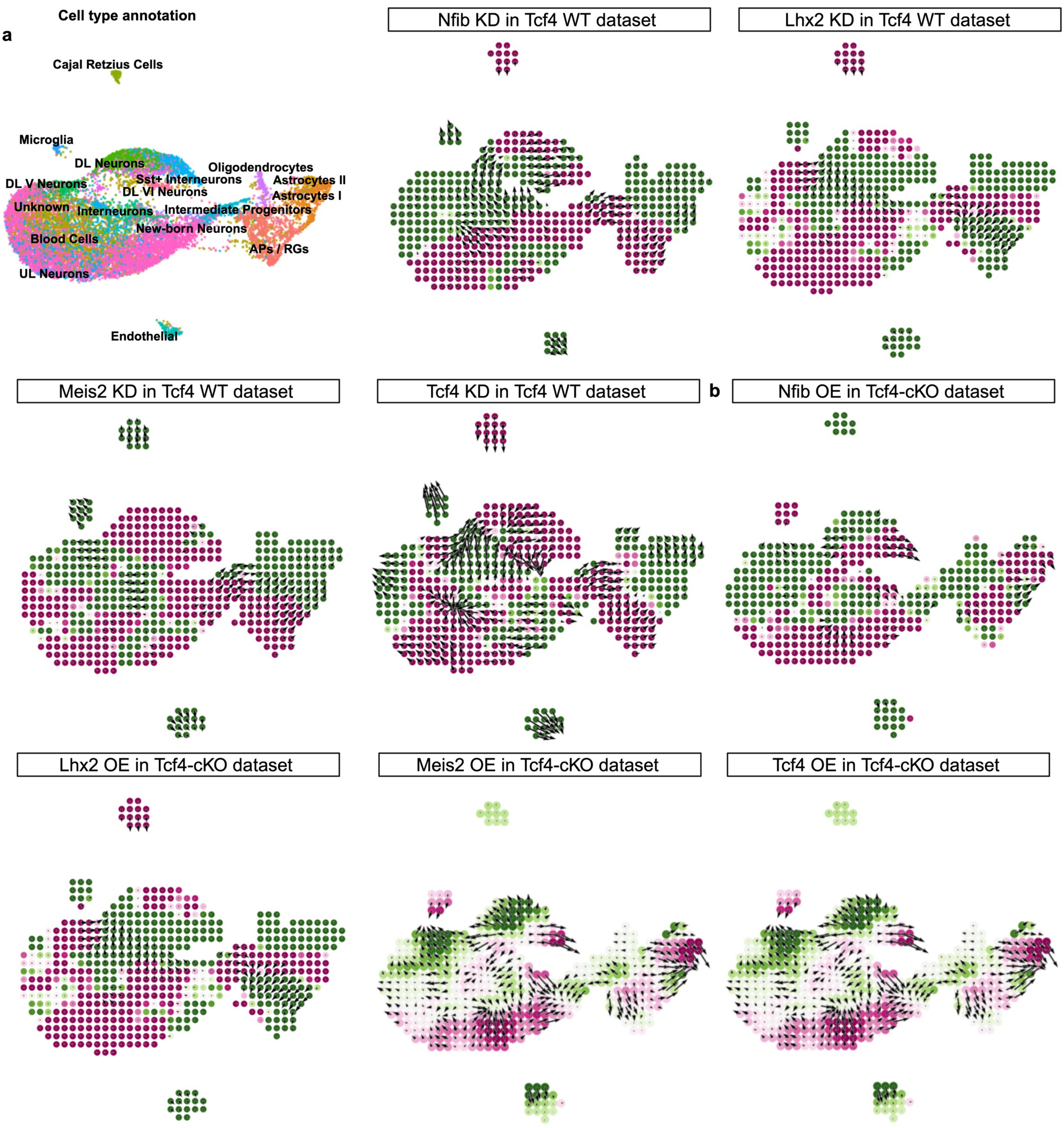


**8. *In silico modelling confirms that Dot1l targeted TFs affect astrogenesis also in a data set from E18.5 WT and Tcf4 LOF cells***

**a** UMAP elucidating the cell types in E18.5 *Tcf4* cKO dataset (left). Colour represents different clusters. Simulation vectors show the predicted transition in cell states after *Nfib, Lhx2, Meis2, Tcf4* KD individually representing the altered likelihood of a cell state change post KD (right). **b** Simulation vectors show the predicted transition in cell states after *Nfib, Lhx2, Meis2, Tcf4* OE individually representing the altered likelihood of a cell state change post OE. Pink and green denote the effects of perturbation on lineage trajectories. Pink indicates a deviation from the expected developmental trajectory, whereas green indicates that the differentiation trajectory remains aligned with the expected developmental lineage. The arrows represent the predicted shift in cell identity in response to the TF perturbation.

### *Supplementary 9*

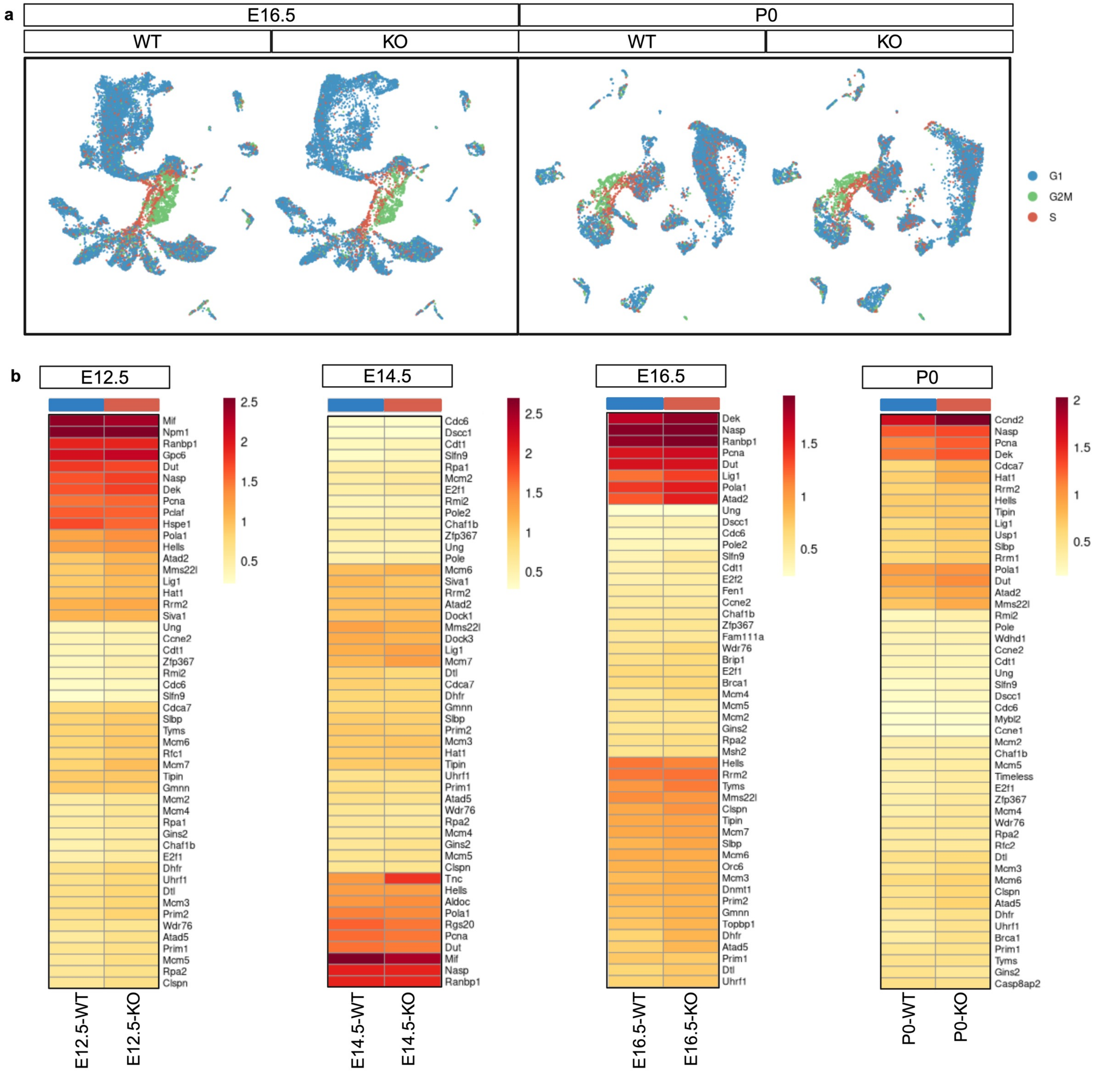


**9.** ***Expression of S-phase genes upon DOT1L LOF is mostly affected at E12.5 compared to later developmental time points***

**a** UMAP highlighting the cell cycle phase (G1, G2M, S) in E16.5 dataset (left) and P0 (right). **b** Heatmap showing the expression of top 50 S-phase genes per timepoint (E12.5-P0) across genotypes (WT: blue, cKO: orange).

### Table S1

| | | Astrogenesis | | | | Conversion | |
|---|---|---|---|---|---|---|---|
| | | APs -> AI | APs -> AII | AI -> APs | AII -> APs | AI -> AII | AII -> AI |
| **P0** | **KO** | Creb5 | Tcf4<br>Nfib<br>Meis2<br>Lhx2 | | | Tcf4<br>Sox6<br>Meis2<br>Nfib<br>Nfia<br>Lhx2 | Creb5 |
| | **OE** | Nfia<br>Sox6<br>Creb5<br>Lhx2 | Lhx2 | | Tcf4<br>Nfib | | Tcf4<br>Nfib<br>Meis2<br>Sox6<br>Creb5 |
| | | APs -> EA | BPs -> EA | EA -> APs | EA -> BPs | | |
| **E16** | **KO** | Meis2 | Meis2<br>Tcf4<br>Lhx2<br>Nfib<br>Emx1 | Lhx2<br>Creb5<br>Nfia<br>Sox6 | | | |
| | **OE** | Emx1<br>Sox6<br>Nfia<br>Creb5<br>Lhx2 | Sox6<br>Nfia<br>Creb5 | Tcf4<br>Meis2<br>Nfib?<br>Creb5 | Tcf4?<br>Creb5 | | |
